# Flagellar energetics from high-resolution imaging of beating patterns in tethered mouse sperm

**DOI:** 10.1101/2020.08.31.269340

**Authors:** Ashwin Nandagiri, Avinash S. Gaikwad, David L. Potter, Reza Nosrati, Julio Soria, Moira K. O’Bryan, Sameer Jadhav, Ranganathan Prabhakar

## Abstract

While much is known about the microstructure of sperm flagella, the mechanisms behind the generation of flagellar beating patterns by the axoneme are still not fully understood. We demonstrate a technique for investigating the energetics of flagella or cilia. We record the planar beating of tethered wildtype and *Crisp2*-knockout mouse sperm at high-speed and high-resolution and extract centerlines using digital image processing techniques. We accurately reconstruct beating waveforms using a Chebyshev-polynomial based Proper Orthogonal Decomposition of the centerline tangent-angle profiles. External hydrodynamic forces and the internal resistance from the passive flagellar material are calculated from the observed kinematics of the beating patterns using a Soft, Internally-Driven Kirchhoff-Rod (SIDKR) model. Energy conservation is employed to further compute the flagellar energetics. We thus obtain the distribution of mechanical power exerted by the dynein motors without any further assumptions about mechanisms regulating axonemal function. We find that, in both the mouse genotypes studied, a large proportion of the mechanical power exerted by the dynein motors is dissipated internally, within the passive structures of the flagellum and by the motors themselves. This internal dissipation is considerably greater than the hydrodynamic dissipation in the aqueous medium outside. The net power input from the dynein motors in sperm from *Crisp2*-knockout mice is significantly smaller than in corresponding wildtype samples. The reduced power is correlated with slower beating and smaller amplitudes. These measurements of flagellar energetics indicate that the ion-channel regulating cysteine-rich secretory proteins (CRISPs) may also be involved in regulating mammalian sperm motility.

## I. INTRODUCTION

In their journey towards the oocyte, sperm propel themselves by beating a whip-like flagellum. This motility is essential for successful fertilization and is fundamental to reproduction. Understanding sperm motility is essential for improving male infertility treatments, animal breeding, and wildlife conservation [1]. Despite the vast body of work on the structure and function of different parts of the axoneme - the internal “engine” powering the flagellum [2–4] – and other accessory structures that surround the axoneme, such as the outer dense fibers [5] and the fibrous sheath [6], the mechanisms that control the complex beating patterns observed in flagella remain poorly understood [7–10]. It is, however, recognized that mechanical properties of the flagellum and its surroundings play a crucial role in determining sperm motility [1]. Measurements of the mechanical behavior of single flagella in living sperm have however remained a critical bottleneck.

We demonstrate here a set of powerful new tools that enable detailed calculation of the mechanical energetics of single sperm flagella from high-resolution optical microscopy. Our approach is based on the idea that the beating pattern carries information about internal dynamics. Automated image-analysis tools have long been used to study sperm movement [11–13]. Computer-aided Sperm Analysis (CASA) systems are today used extensively in clinical settings to rapidly assess the viability of samples containing hundreds of cells in a single field of view [14]. These high-throughput techniques, however, do not resolve flagellar motion. Improvements in digital imaging and storage have now placed within reach the highspeed, high-resolution and long-exposure imaging that researchers of flagellar propulsion have long sought [15–18]. A wide range of digital image processing algorithms are now available [19] that can be combined with high-performance parallel computing to analyze thousands of video frames with little manual intervention [20–24]. We have implemented these image-analysis techniques to automatically extract centerlines of sperm flagella in every video frame.

To quantitatively analyze beat patterns in a statistically meaningful way, we need to image swimming sperm over several beat cycles. While rapid progress is being made on full three-dimensional tracking [25–27], it is unlikely that sufficient beat cycles can be reliably recorded with freely swimming sperm that can quickly move out of focal plane or the field of view [28]. Instead, we image flagella beating freely in the focal plane in cells tethered chemically at their heads to a cover-slip. Our tethered-cell assay, in principle, permits imaging single cells until they stop beating. We report here results obtained by analyzing large numbers of (~ 50) beat cycles in single tethered sperm in freshly prepared samples when they are most vigorous [29].

Beating patterns in sperm flagella have been studied previously to investigate changes induced by environmental factors [22, 30; 31] or by gene mutations [32; 33]. We build here on the suggestion that the technique of Proper Orthogonal Decomposition (POD) can be applied on the time-resolved tangent-angle profiles of flagellar centerlines to analyze their kinematics [22; 34; 35]. POD is widely applied in the analysis of turbulent flows [36; 37] and other fields [38; 39] to reduce complexity of spatiotemporal patterns and represent them with a much smaller set of numbers, while still retaining accuracy. To objectively compare flagellar beating patterns, we apply POD to unambiguously identify the *mean beat cycle* of each sperm from the time-series of the tangent-angle profiles of its flagellar centerline. We can compute average cycles of any kinematic or dynamic quantity derived from the tangent-angle profiles. We further introduce a technique to consistently represent the POD shape modes with smooth Chebyshev polynomials to ensure that the tangent-angle profile is sufficiently smooth and its spatial derivatives can be computed without spurious artefacts. The tangent-angle profile obtained thus is consistent with the rigid-body kinematics of the stiff head region. This Chebyshev-POD (C-POD) technique allows for efficient calculation of geometric quantities such as the local curvature and kinematic quantities such as the velocity components, at any material point on the centerline.

We use this geometric and kinematic information to determine the hydrodynamic resistance offered by the external fluid medium using Resistive Force Theory [40; 41], and further calculate internal forces by applying conservation principles. This requires a model for the mechanical behavior for the flagellar body. Several models have been proposed that consider the flagellum to be an “active” material [24; 42–46]. These are based on different models for motor forcing in the axoneme and the regulation of their kinetics. We propose instead a different approach that is agnostic to the nature of motor activity and avoids invoking the assumption that the flagellar material is active. We consider the motion of the non-motor passive material of the flagellum under the action of the unknown forces exerted by the axonemal motors. This allows us to use well-established principles for the continuum material stress in the passive flagellar material. The resulting Soft, Internally-Driven Kirchhoff Rod (SIDKR) model leads to an energy balance across the flagellum, which we then use to determine the spatio-temporal distribution of motor power across the flagellum over its mean cycle.

We have used this approach to analyze flagellar beating patterns of sperm from wildtype (WT) and *Crisp2* knockout (KO) mice. The CRISPs (cysteine-rich secretory proteins) are a group of proteins which are predominantly expressed in the male reproductive tract [47]. CRISP2 is incorporated into the sperm acrosome, connecting piece and the outer dense fibers of the sperm tail. It is known that the deletion of *Crisp2* in mice leads to compromised sperm function, including altered sperm motility [33; 48]. The precise effect on flagellar function, however, is unknown. Our observations with these sperm reveal intriguing new information: there is considerable intracellular friction within the flagellum. This challenges the widely held view that the hydrodynamic resistance offered by the viscous fluid medium outside is the sole dissipative sink that must be overcome by the continual driving provided by the dynein motors. Further, the flagellar filament is also conventionally regarded as an elastic body that perfectly stores energy temporarily by bending. Our findings suggest instead that internal friction within the passive structures of the flagellum, and within the motors themselves, may be as large as the external hydrodynamic friction. These are in line with recent observations also made in algal cilia [28]. These sources of internal dissipation could therefore play a significant role in determining beating patterns in sperm [43]. This insight could be vital for understanding dramatic changes in flagellar beating patterns induced by changes in the medium [31] or the proximity of surfaces [49; 50].

### A. Theoretical model

#### 1. The Soft, Internally-Driven Kirchhoff Rod Model

Flagellar motion is driven internally by the action of dynein motors distributed within the axoneme. The sperm body is treated as a slender, flexible filament immersed in a viscous fluid (Figure 1). It is assumed that the passive material of the sperm body is a Kirchhoff rod [51–53], *i.e.*, it is inextensible and each of its material cross-sections remain rigid and planar, while rotating with respect to each other about the rod axis as it bends and twists. The passive Kirchhoff rod has external as well as internal surfaces. It is driven by axonemal motors acting on its internal surfaces and the resulting motion is resisted by the hydrodynamic forces that act on its external surface (Figure 1) as well as the stresses that arise to resist material deformation as the rod bends.

**FIG. 1.**
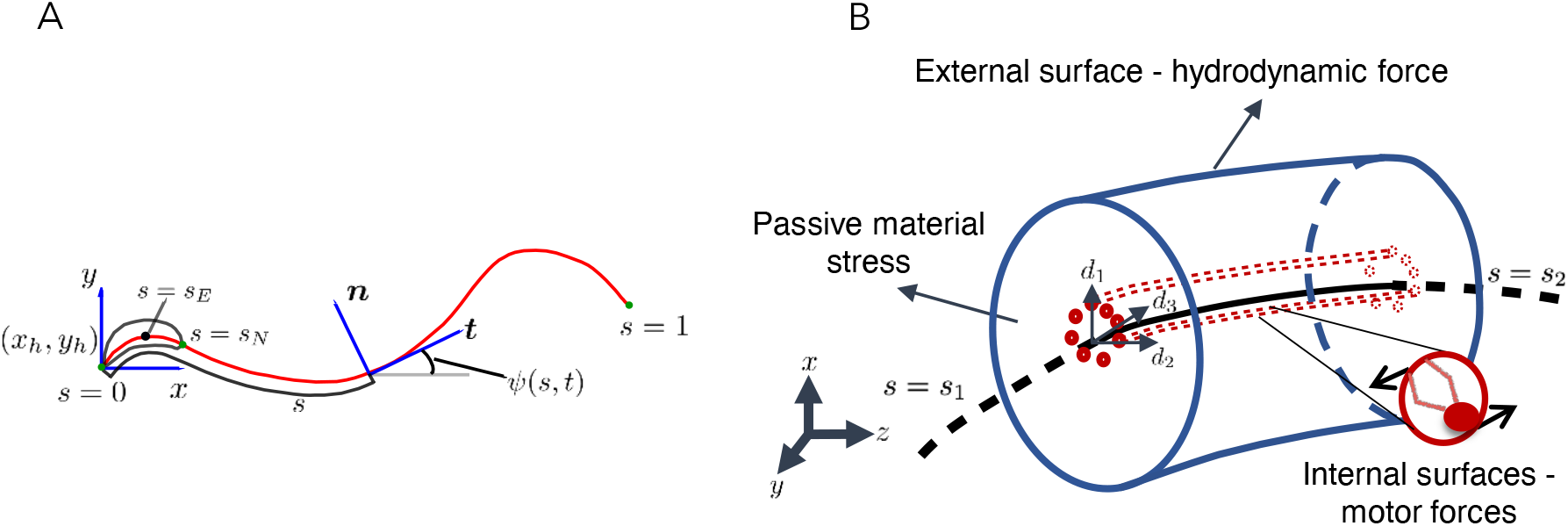
Schematic representations for the Soft Internally-Driven Kirchhoff Rod model: (A) Geometric variables defined along the centerline. (B) An arbitrary control volume used for deriving the equations of the model: the volume consists of the passive flagellar material; hydrodynamic forces act on the external surface while axonemal motors act on the internal surfaces. The passive material adjacent to the cross-sectional faces at either end exert stresses on those faces.

The instantaneous space curve of the axial centerline of the filament, (*s, t*), is parameterized by its arc length variable, *s*, defined such that *s* = 0 at the tip of the head, and *s = L* at the tail end. A local material frame is attached to each cross-sectional plane, and is specified by a triad of unit vectors, d_*k*_, where *k* = 1, 2, 3. In general, the smooth variation of these vectors with *s* at any instant of time, *t*, is specified in terms of the Darboux vector, Ω, where *∂*d_*k*_/*∂s* = Ω × d_*k*_. The components Ω_*k*_ of the Darboux vector are the generalized curvatures. Since we shall only consider motion of the rod in the *x − y* plane, we align the material frame at each cross-section with the Frenet-Serret frame associated with each point on the axial curve. For this choice, d_1_ = t = *∂*r/*∂*s, the unit tangent vector to axial curve. The other two vectors, d_2_ = n, and d_3_ = b, are the normal and binormal vectors, which span the cross-sectional plane. The Darboux vector for the Frenet-Serret frame is Ω = *T*(*s, t*)d_1_ + *C*(*s, t*)d_3_, where *C* and *T* are the curvature and torsion profiles at any time. For planar motion, b = e_*z*_ (pointing out of the plane of the page) is a constant; hence, *T* = 0. The geometry of a planar Kirchhoff rod at any instant is thus fully specified by the curvature, *C*. The velocity of a point on the centerline, v(*s, t*) = *∂*r/*∂t*. Cross-sectional planes can rotate relative to each other. Then, *∂*d_k_/*∂t = ω* × d_*k*_, where *ω*(*s, t*) is the instantaneous angular velocity of a cross-sectional plane at *s*. It can further be shown that *ω* and Ω are related through,

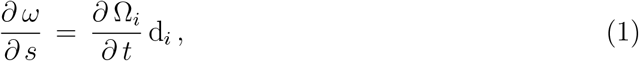

where Einstein’s summation convention is used. This implies that, for planar motion, where *ω = ω* e_*z*_,

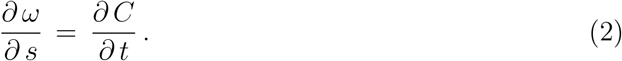

For inertialess rods, consideration of the conservation of linear momentum for a segment of the rod where *s* ∈ [*s*_1_, *s*_2_] formally yields the following equation (see Supplementary Information; [54]):

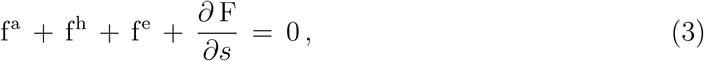

where f^a^(*s, t*) and f^h^(*s, t*) are the force distributions per unit length on the cross-section at any s due to the surface tractions exerted by internal motor activity and the external hydrodynamic resistance, respectively. Other external forces, such as the force exerted by a tethering traction at a wall, are accounted for by the distribution f^e^(*s, t*). The passive stress in the Kirchhoff rod results in a force, F, exerted on a cross-section by the material on its aft side. The gradient with respect to s of F in the momentum balance thus describes the *net* restoring force per unit length on a cross-section due to passive internal stresses resisting deformation.

From conservation of angular momentum, we obtain [54]:

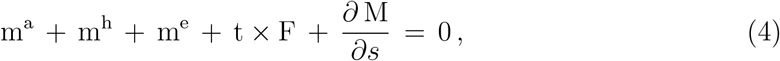

where m^a^(*s, t*) and m^h^(*s, t*) are the torques per unit length exerted by the surface tractions due to the internal motors and the external viscous hydrodynamic resistance; m^e^ is the torque distribution due to other external forces. The torque on a cross-section exerted by the passive material stresses on its aft side is M, and its gradient in the equation above is the net restoring torque distribution.

Energy conservation further shows that at any cross-section, in general,

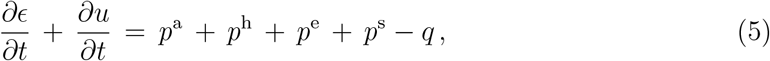

where *ϵ*(*s, t*) is the local elastic energy per unit length of the rod and *u*(*s, t*) is the thermal internal energy distribution. On the right-hand side, *q* is the net rate of heat removal per unit length of the rod by the surroundings, while each of the remaining terms is, respectively, the mechanical power per unit length delivered into the rod cross-section by the action of the motors, the hydrodynamic and non-hydrodynamic external forces, and the passive material stress. The motor power distribution, *p*^a^, is the key unknown in our study. The hydrodynamic power distribution is related to the corresponding force and torques distributions:

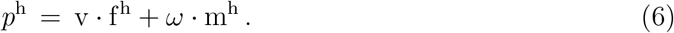

The other external mechanical power *p*^e^ is similarly related to the external force and moment distributions, f^e^ and m^e^. The net rate of work done on a cross-section by the action of the local stress gradient is

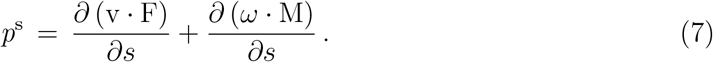

The sign convention used here is that mechanical power due to work done *on* a cross-section of the rod and tending to increase the local internal energy storage is positive whereas the power due to work done by that cross-section to overcome resistances leading to a decrease in stored energy is negative. Due to its purely dissipative nature, *p*^h^ is therefore always negative at any *s* and *t*. In our study, the external force and moment due to the tethering constraint exerted on the head cannot be measured directly. The mechanics of this tether could be complex and, at any instant of time, *p*^e^ may be positive or negative. However, over a full cycle, we expect net work to be done by the cell against the tethering constraint.

The key advantage in treating the motor contribution as a forcing that is external to the passive material of the Kirchhoff rod is that we can treat the active forcing as an unknown to be extracted from experimental data in a model agnostic manner while applying well-established concepts to treat passive material stresses within the Kirchhoff rod. The passive stress tensor can be formally split into an elastic and a dissipative part, so that the total material torque, M = M^E^ + M^D^. It can be shown that Equation 5 is satisfied when the elastic torque arising from the passive material stress is such that

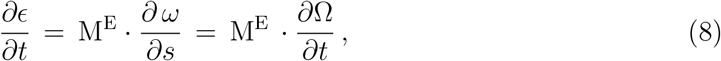

and the dissipative part of the material stress is such that

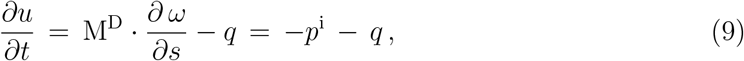

where *p*^i^ denotes the rate of internal frictional dissipation per unit length. Since the material of the Kirchhoff rod is passive, the Second Law of Thermodynamics requires that *p*^i^ ≤ 0 everywhere (with the sign-convention used here) [55]. Since the dynein motors are excluded from the control volume in the analysis above, *p*^1^ does not include any dissipation that occurs within the motors themselves. We shall later discuss how we separately obtain the motor dissipation.

#### 2. Constitutive relations

Although presented in the context of a sperm body, the equations above are generally valid of any inertialess, internally-driven Kirchhoff rod. To proceed further, we make several constitutive assumptions that are specific to the case of a sperm cell tethered at its head. The sperm body is assumed to be composed of a head region, *s* ∈ [0, *s*_N_], and a flagellar tail region, *s* ∈ (*s*_N_, *L*], with *s*_N_ denoting the location of the neck junction between the two regions. We assume that the head is a rigid body. In our experiments, cells are further tethered at a point in the head region, and the head can rotate rigidly about this tether point. Therefore, although the angular velocity *ω* = 0 in the head region, rigid-body kinematics dictates that *∂ω/∂s* = 0 everywhere in the head region. Hence, from Equation 1, *∂*Ω_*k*_/*∂t* = 0 across the head. Therefore, for planar beating, *∂ω/∂s* = *∂C/∂t* = 0 across the head. The flagellar tail is flexible and not subject to the kinematic constraints above.

The head does not contain internal motors, which are all distributed only along the tail region. Therefore, f^a^, m^a^ and p^a^ are all zero for *s* ∈ [0, *s*_N_]. In the flagellar tail, each dynein motor is assumed to act on the internal surfaces of a cross-section such that the forces exerted at its two ends are of equal magnitude but in opposite directions. Therefore, f^a^ = 0. However, the net torque they exert is not zero, and therefore m^a^ ≠ 0, which serves to drive the filament’s motion.

The external hydrodynamic force distribution is given by Resistive Force Theory [40; 41]:

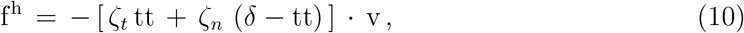

where the tangential and normal hydrodynamic friction coefficients in a fluid medium of viscosity, *μ*, are *ζ_t_* = 2*πμ*/ln(2*L/a*) and *ζ_n_* = 4*πμ*/[ln(2*L/a*) + 1/2], respectively. The corresponding contribution to the hydrodynamic power distribution, v · f^h^ is thus always negative and consistent with the dissipative nature of the hydrodynamic resistance. The cross-sectional radius of the cylindrical filament, *a*, is further not constant along the sperm body. For our calculations here, only the variation of the radius in the tail region is relevant. We assume a linear taper along the flagellum *i.e*. for *s > s*_N_,

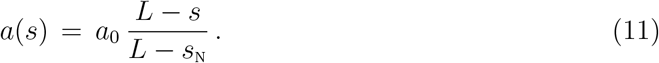

where *a*_0_ is the radius at the neck.

The rigid head region requires no further constitutive assumptions. The tail region can deform and therefore requires a constitutive model that relates its material stresses to its deformation. The simplest constitutive model for the elastic stress in a passive material is the Hookean model which leads to a linear relation between the elastic material torque and the local curvature. The corresponding elastic energy distribution must be consistent with Equation 8. Thus, in the tail region,

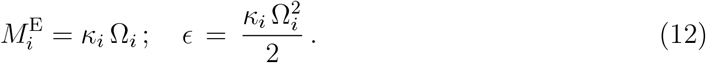

where *κ_i_* is an elastic stiffness coefficient. The simplest constitutive model for the dissipative stress that satisfies the condition imposed by the Second Law that the dissipation rate is positive always leads to the following expression for the dissipative part of the internal torque:

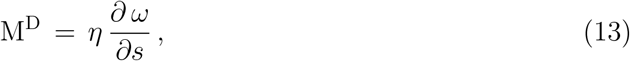

where *η* > 0 is the internal friction coefficient per unit length. Taken together, the constitutive equations above are equivalent to modeling the Kirchhoff rod as a passive viscoelastic Kelvin-Voigt solid [56]. For the linear taper assumed in the tail region, the elastic stiffness and internal friction coefficients can be shown to vary with the radius as *a*^4^ *i.e*.

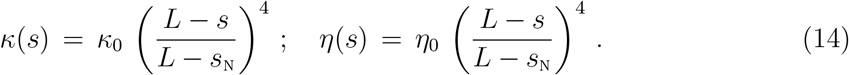

where *κ*_0_ and *η*_0_ are the elastic stiffness and frictional coefficients at the neck. For planar motion, the relations above reduce to *M*^E^ = *κC* and *ϵ = κC*^2^/2. The distributions of the elastic storage rate and power dissipated due to internal friction are, respectively,

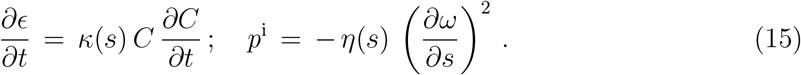

In what follows, we denote the elastic storage rate as 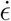.

We make a few other simplifying assumptions. The head and tail ends are free; F and M are, therefore, zero at the two ends. The external surface traction due to tethering at the wall acts at a single location, s_E_, on the head and is zero elsewhere i.e. f^e^ = F^e^*δ*(*s − s*_E_) and m^e^ = M^e^*δ*(*s − s*_E_). The power distribution, *p*^e^, is therefore zero at all points in the tail. The system is further isothermal and changes in the internal thermal energy of the body are negligible *i.e. ∂u/∂t* = 0. Any internal frictional heat generation is, therefore, instantaneously balanced by the heat removal to the surroundings, *q*. As far as the energetics of the flagellum is concerned, the ratio of the contributions from the external hydrodynamic moment, m^h^, and the hydrodynamic force, f^h^, to the total hydrodynamic power i.e. the ratio |*ω* · m^h^| / |v · f^h^| is expected to scale as *a/L* ≪ 1. The contribution of m^h^ to flagellar energetics is therefore neglected.

Using the relations above, Equation 5 can be rearranged to yield the active power distribution for the tail region:

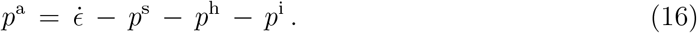

As we show in, all the terms on the right-hand side can be determined from centerline kinematics, and therefore, the active power distribution can be obtained from analysis of experimental videos of flagellar beating in sperm. The total instantaneous storage rate, *Ė*, and the hydrodynamic, passive internal frictional, and active powers *viz. P*^h^, *P*^i^ and *P*^a^, are calculated in the tail region by respectively integrating the distributions 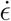, *p*^h^, *p*^1^ and *p*^a^ over the tail. We can similarly obtain the powers over just the mid-piece or over the principal-piece alone.

The input parameters required for these calculations are the medium viscosity, *μ*, the body length, *L*, the neck radius *a*_0_, and stiffness and friction coefficients at the neck, *κ*_0_, and *η*_0_. Other than the last two, the remaining parameters are directly measurable from experiments. From the few measurements of the bending stiffness in the literature, we use *κ*_0_ = 7 × 10^4^ Pa *μ*m^4^in calculations here for mouse sperm [9]. There are no clear measurements yet of *η*_0_, however. Since *η*_0_/κ_0_ represents a characteristic viscoelastic timescale for the passive flagellar material, if internal viscoelasticity plays an important role in flagellar dynamics, one would expect 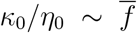, the observed mean frequency of a beat cycle. This leads to a scaling estimate of 10^3^ Pa s *μ*m^4^for *η*_0_. We first report below the results obtained for flagellar energetics with the scaling estimate and subsequently examine the qualitative effect of the choice of *η*_0_ on these results.

#### 3. Energetics from image-analysis and POD

In, we describe the image-analysis and data-processing algorithms used to obtain power distributions from microscope videos of tethered sperm samples from WT and *Crisp2* KO mice. The image-analysis algorithm is used to process videos of single sperm cells tethered to a glass surface and beating in the focal plane of the microscope and extract centerlines of sperm bodies in every video frame. 20th-order Chebyshev polynomials are fitted through these centerlines to construct tangent-angle profiles (see Figure 1(A)) of the flagella. The Chebyshev polynomials are designed to be consistent with the rigid-body kinematics of the head region.

In general, the POD is an order-reduction technique that optimally approximates spatiotemporally varying data. In our C-POD approach, we apply POD on the time-dependent Chebyshev coefficients to represent the deviation of *ψ*(*s, t*), the time-resolved tangent-angle profile of the centerline from its time-average, ≠_0_(*s*), as a weighted sum of *M* orthogonal shape modes (see). In other words,

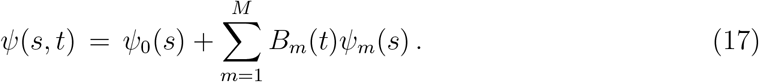

The set of “shape modes”, *ψ_m_, m* = 1… *M*, is optimal in the sense that, for any given *M*, the approximation *ψ* is guaranteed to deviate least from the original data than any other expansion in terms of another set of M mutually orthogonal basis functions [35; 37]. We describe in the C-POD method to obtain shape modes that are each a 20-th order, Chebyshev polynomial that is consistent with the head region executing rigid-body rotation. The corresponding time-dependent weights of the shape modes are referred to as “shape coefficients”.

With the smooth C-POD tangent-profiles, we can efficiently compute at any *s* and *t*, geometric and kinematic quantities in the beating plane, such as the curvature *C* and its derivatives with respect to *s* or *t*, the flagellar velocity v and the cross-sectional angular rotation rate, *ω*. The algorithm for the calculation of dynamic quantities such as force, torque and power distributions from the kinematics of the flagellum is described in.

## II. RESULTS

### A. POD enables identification of beat cycles

Figure 2 summarizes generic observations on the C-POD shape-modes and their coefficients. In all the results presented here, the arc-length coordinate *s* along the centerline is normalized by the maximum observable length of the whole flagellum in the entire duration of a sample video. The mid-piece region corresponds approximately to values of *s* in the range 0.1 to 0.3, and the principal piece extends from *s* = 0.3 to *s* = 0.85.

**FIG. 2.**
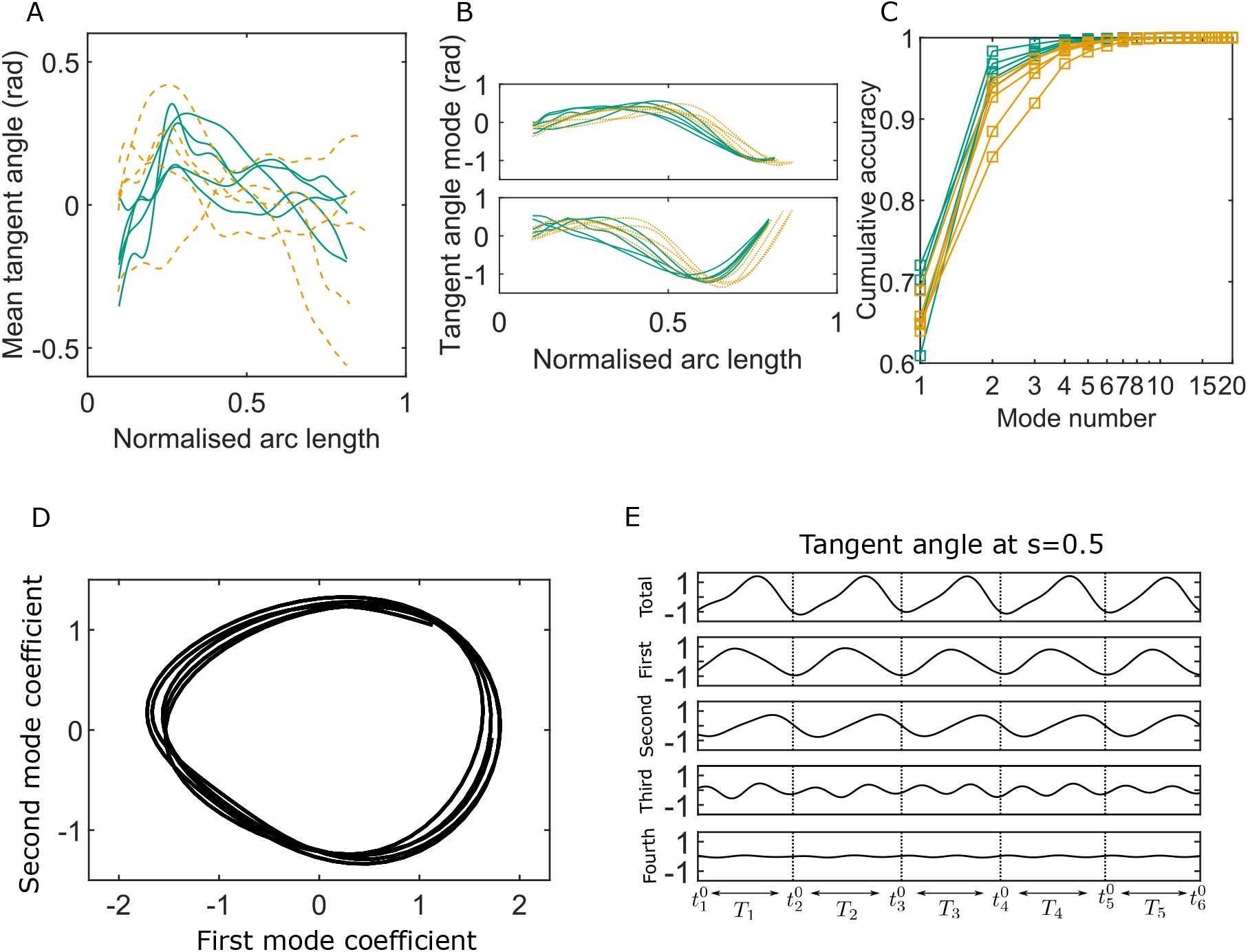
Key variables of the Chebyshev-polynomial-based Proper Orthogonal Decomposition (C-POD) of experimental tangent-angle data. (A) Time-averaged tangent-angle profiles for five WT (continuous curves) and five KO (dashed curves) samples. (B) First (top) and second (bottom) C-POD shape modes for WT (continuous curves) and KO (dashed curves) samples; the colors are as in A. (C) Cumulative accuracy of the C-POD representation for WT and KO samples; the colors are as in A. A representation using the first four modes captures 95% or more of the observed centerline shapes for all samples. (D) Five shape cycles for a single WT sample in the parameter-space defined by the time-dependent coefficients of the first two C-POD shape modes. The zero-crossing of the second modal coefficient marks the start of a new cycle. (E) Contributions of the first four modes to the tangent angle at the mid-point of the sperm body in the five tangent angle cycles in D: the horizontal line in the top plot is the time-averaged tangent angle at for this WT sample. The starting time of the i-th cycle is denoted as 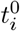 and its duration (i.e. cycle time) is *T_i_*.

Mouse sperm have distinctive hook shapes. In the image-processing protocol we have followed, all video frames are initially rotated such that the head is on the left end of the body with the hook facing concave-downward. For most of the WT and KO samples, the time-averaged tangent angles (*≠*_0_(*s*)) are observed in Figure 2 (A) to consistently first increase with *s* around the mid-piece region, before decreasing in the principal piece. Since the local curvature, C = ∂≠/∂s, the gradient of the tangent angle with respect to s, Figure 2 (A) shows that the time-averaged shape for these samples is curved such that it is concave in the anti-hook direction in the mid-piece and concave in the pro-hook direction in the principal piece. The mean shapes thus show that the asymmetric spatial bias in the beating pattern over time is not uniform across the flagellum. In the one outlier KO sample (KO-5) in Figure 2 (A), however, the mean shape is anti-hook concave throughout.

The periodic beating of the flagellum about the mean shape is described by the C-POD shape modes and their time-dependent coefficients. The shapes of the first two shape modes (*≠*_1_(*s*) and *≠*_2_(*s*)) in Figure 2 (B) are qualitatively similar across the WT and KO samples. The key advantage of using the POD method to represent beating patterns is its optimality: a significant proportion of the beating pattern can be studied and understood by considering just a few shape modes. Figure 2 (C) plots the the cumulative contribution of the shape modes to the overall accuracy in capturing the full centerline datasets. Just the first two modes achieve a capture efficiency greater that 92% for all the WT samples, and for three out of the five KO samples. Even for the other two KO samples, these dominant shape modes account for more than 85% of the observed beating patterns. Across all samples, the first four modes describe at least 95% of the beating patterns. We therefore calculate all kinematic, dynamic and energetic quantities using the first four shape modes and their time-dependent coefficients.

As pointed out by Werner *et al*. [35] and Ma *et al*. [34], the periodicity in the beating pattern is clearly brought out by plotting the coefficients *B*_1_(*t*) and *B*_2_(*t*) of the two dominant modes against one another. For any sperm sample, the trajectory traced out in *B*_1_-*B*_2_ phasespace consists of loops, one for each beat cycle (*e.g*. Figure 2 (D)). We can clearly demarcate the start and end time for each beat cycle as the time at which the polar angle in the *B*_1_-*B*_2_ phase-space crosses zero. Thus, the overall time-series for any quantity can be split into individual beat cycles, as demonstrated Figure 2 (E). This also means that in each sperm sample, the shape at the start of a beat cycle always corresponds mostly the shape of the first dominant mode (with minor contributions from modes higher than the second; Figure 2 (E)). Although Figure 2 (D) and (E) show only a few cycles for clarity, the C-POD technique applied to tethered sperm, makes it possible to systematically accumulate data for large numbers of beat cycles and quantitatively compare, in a statistically meaningful sense, individual sperm samples within a genotypical population, and also compare one genotypical population with another.

### B. Active power distribution provides evidence for energy dissipation by dynein motors

Before comparing the behaviour of the WT and KO samples, we first present the energetic flows typically observed in any sample. [Figure 3 plots the kymographs for the different energetic contributions obtained with the scaling estimate of the internal friction coefficient, *η*_0_ = 10^3^ Pa s *μ*m^4^, over several beat cycles for one of the WT samples. Similar results are obtained for all the other samples. The banded structures in these kymographs provide a visual confirmation of the spatiotemporal periodicity of the energy flows corresponding to the periodic beating of the flagellum.

**FIG. 3.**
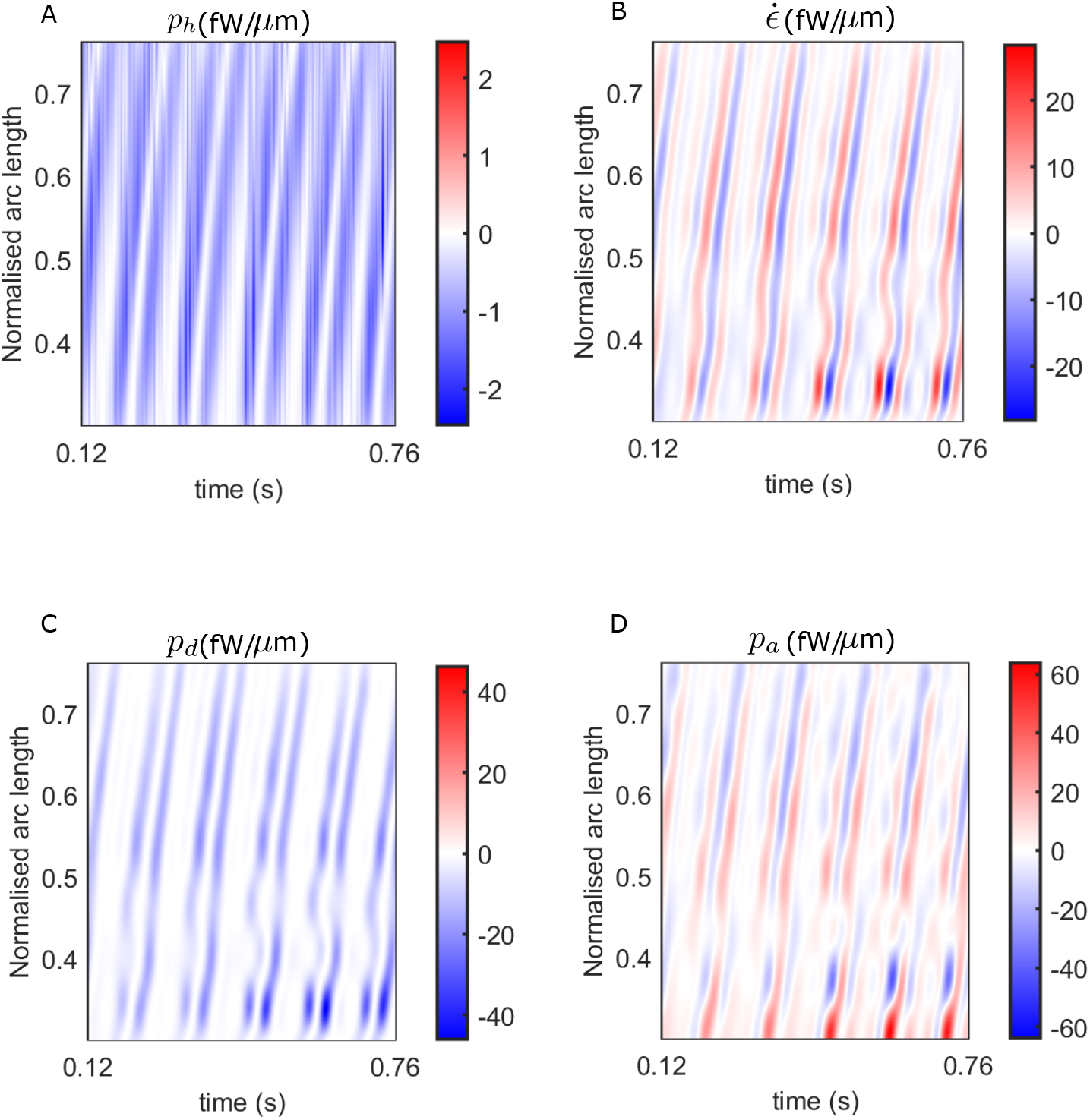
Spatiotemporal distributions of the rates of energetic quantities along the flagellum of a WT sperm over several beat cycles: red indicates positive rates, while blue indicates negative rates. The data in C and D have been obtained using the scaling value of 10^3^ Pa s m for the internal friction coefficient.

In Figure 3 (A), the hydrodynamic power distribution, *p*^h^, is always negative: that is, every part of the flagellum is at all times working against the hydrodynamic forces exerted externally by the viscous environment provided by the ambient fluid. This work done on the fluid is dissipated away by fluid friction. The elastic storage rate per unit length, 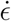 at any location s, however, alternates between positive (red) and negative (blue) values in Figure 3 (B). As a bending wave propagates through that location, the local curvature at that *s* increases, leading to potential energy being stored elastically and a positive rate of 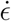 at that location. As the filament begins to relax and straighten out, the stored elastic energy is released and begins decreasing leading to negative 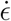 values there. The filament then proceeds to bend in the other direction at that point, leading to a second positive growth of 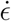 within the same beat cycle, followed by a negative phase in 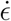 as the filament relaxes back towards being undeformed and straight at that location. Thus, at any s in Figure 3 (B), each beat cycle consists of two successive positive and negative growth-rate phases in 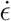.

Comparing the bands in Figure 3 (B) with those in (A), it is clear that every single planar wave that propagates down the filament is associated with a pair of hydrodynamic dissipation peaks: the contribution of any single location to the hydrodynamic dissipation peaks as the filament moves quickly while bending and relaxing back on one side, and then again, on the other side. These bands are mirrored in Figure 3 (C), which plots *p*^i^, the distribution of power dissipated due to *internal* friction. This frictional dissipation, calculated with *η*_0_ = 10^3^ Pa s *μ*m^4^, is due to relative motion between adjacent cross-sectional planes of the flagellar material, which also peaks at a location when a bend towards one side or the other propagates past that point.

The external and internal dissipations and temporary elastic storage of energy must together be supported by the mechanical power input provided by the dynein motors acting on the microtubule surfaces of the flagellum. Figure 3 (D) plots the distribution of this active power input, *p*^a^, across the filament. Surprisingly, we find that the *p*^a^ distribution displays clear negative bands that repeatedly occur in all beating periods and are spread throughout the filament. The positive domains (red) of the *p*^a^ kymograph in Figure 3 (D) represent ATP free-energy being delivered as mechanical power by the motors. In those regions, the motors cause relative sliding of microtubule doublets to rotate the local crosssectional planes in the same sense as the torques they exert *i.e*. since *p*^a^ = *ω · m*^a^, *p*^a^ is positive at a cross-section when both the rotational velocity of that plane, *ω*, and the torque per unit length, *m*^a^, exerted by the dynein motors in that plane have the same sign. On the other hand, where *p*^a^ is negative (blue) in Figure 3 (D), *ω* and *m*^a^ are opposite in sign. At any such point, work is being done *by* the rest of the flagellar material *on* the axonemal motors, driving them back *against* the torque they continue to exert. Experiments with optical tweezers have shown that mechanical work done can be done *on* dynein motors to move them, either in the presence or absence of ATP in the surrounding medium [57]. This energy transferred mechanically back to the motors can neither be stored either within the dyneins nor converted back to chemical free energy (i.e. ATP): it must be therefore quickly dissipated locally within the axoneme itself. This axonemal *motor dissipation*, measured by the negative domains of *p*^a^, is a second source of dissipation within the flagellum, quite distinct from the dissipation, *p*^i^, that is due to internal friction arising from the relative motions of all the other structures in the flagellum that surround the axonemal motors, such as the microtubules, the outer dense fibers, *etc*. By adding together the *p*^a^ distribution over all the locations where it is negative, we can calculate, *P*^m^, the instantaneous rate of energy dissipation due to the dynein motors themselves. The sum of *P*^m^ and *P*^i^ is the total mechanical power dissipated within the whole flagellum.

### C. Means and distributions of beating patterns and energetic variables can be obtained

The qualitative features of the distributions of the key energetic variables discussed above are common to both WT and KO samples. Before identifying significant differences between the beating patterns and energetics of the genotypes, it is worth examining the sample-to-sample variability within each population. Figure 4 (A) shows the mean cycle of the beating pattern in physical *x-y* space for each sperm sample in our study. Flagellar centerlines at the same value of the fractional duration of the mean beat cycle have the same color in Figure 4 (A). This fractional duration of the mean cycle is referred to as the time-phase and is denoted as *τ*. To obtain the mean centerline shape at a particular value of *τ*, we collect, at that *τ*, the *x* and *y* coordinates obtained (using Equation 37) for all the beat cycles, and then calculate their mean values. The bands in Figure 4 (A) around the mean centerlines are the standard errors in the mean (SEM) *y* coordinates at each *s*. Our procedure for identifying the start and end of each beat cycle thus enables calculation and comparison of average beating patterns.

**FIG. 4.**
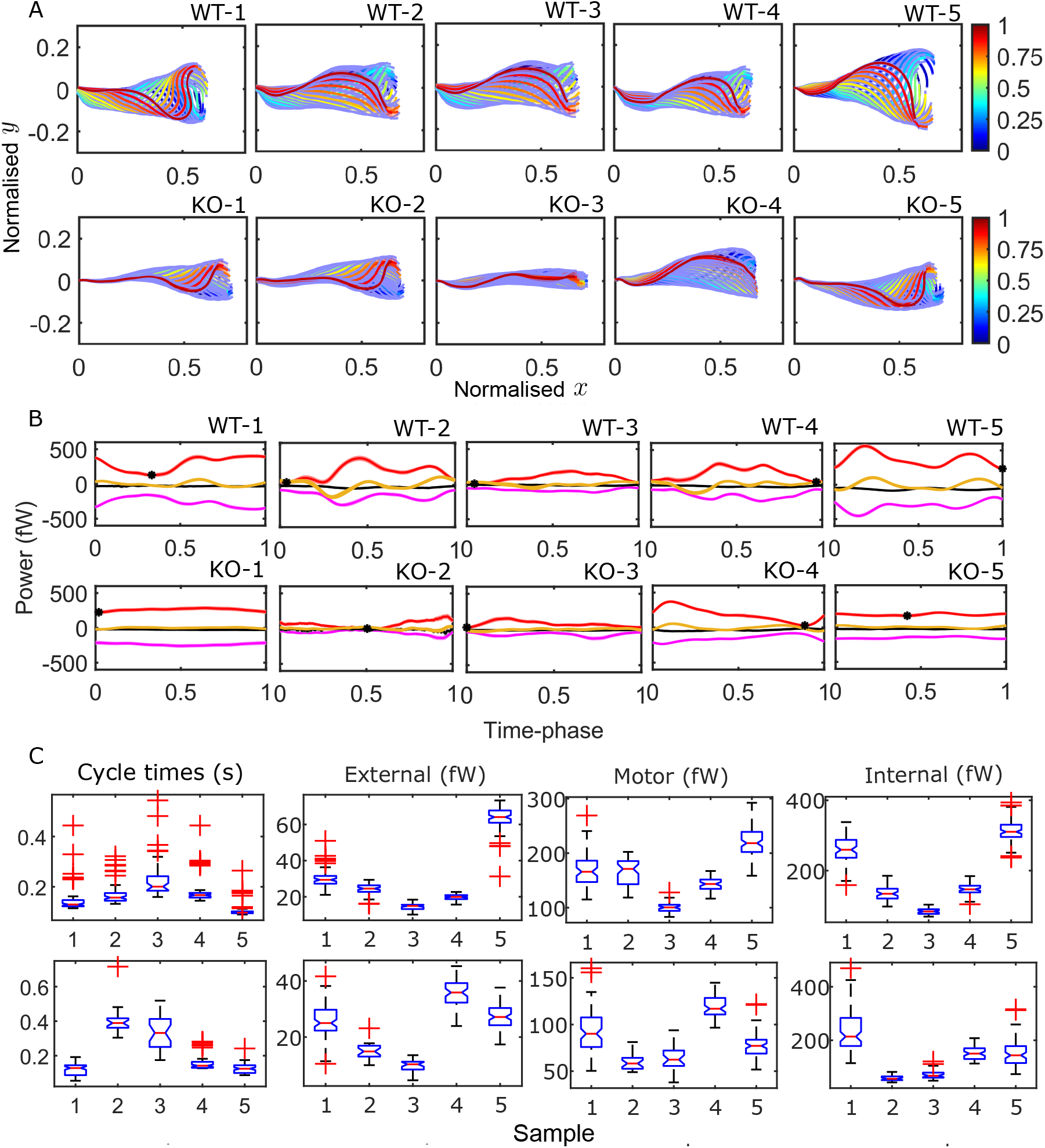
Mean cycles of beat patterns and energetics. (A) Each colored curve shows the mean shape at a particular phase of the mean cycle for the five WT (top row) and KO (bottom row) samples. The color bands around each curve indicates the standard error in the mean-component. (B) Mean cycles for the magnitudes of the net elastic storage (yellow), hydrodynamic dissipation (black), internal frictional dissipation (magenta) and active power (red) in WT (top panel) and KO (bottom panel) sperm samples corresponding to those in A. Bands show standard errors in means. (C) Statistical distributions of cycle times and dissipation rates in each of the WT (top panel) and KO (bottom panel) samples. The box-plots present the median (red line), the first and third quartile (bottom and top box edges), and minimum and maximum (lower and upper whiskers) cycle, when the first shape mode is dominant, suggesting that the first shape mode could be associated with a state of minimum power input. In two of the KO samples (KO-1 and KO-5), however, the net motor power remains nearly constant over the entire cycle.

The difference between the mean beat patterns of the WT and CRISP2 KO samples is striking. The KO samples exhibit a smaller amplitude across the entire flagellar tail. We have quantitatively analyzed the differences in beating patterns caused by mutations in genes encoding for CRISP2 and other CRISPs in a separate study [47]. In Figure 4 (B) and (C), we apply the idea of calculating mean cycles to the energetic variables calculated from the 4-mode C-POD of the tangent-angle profiles. Figure 4 (B) compares the mean cycles in the net rates of elastic storage, (*Ė* yellow), hydrodynamic (*P*^h^; black) and internal frictional (*P*^i^; magenta) dissipations, and the net rate of motor power input (*P*^a^; red curves). At each time-phase, *τ*, in a beat cycle, these mean rates are calculated by collecting the values of *Ė P*^h^, *P*^i^ and *P*^a^ from all the cycles and averaging those values. No distinctive cyclic patterns are immediately apparent in Figure 4 (B). In seven of the ten samples, the minimum value in *P*^a^ (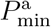; black symbols in Figure 4 (B)) occurs close to the beginning or end of the It is visually apparent from Figure 4 (B) that the mean cycles of the energy flows vary considerably from sample to sample. We plot the distributions of cycle times for each of the WT (Figure 4 (C); top panel - i) and KO (bottom panel - i) samples. Also shown as box-plots are the statistical distributions of the magnitudes of the cycle hydrodynamic, passive internal friction and motor dissipation powers. The cycle power in any single cycle are calculated by integrating an instantaneous power with respect to time over that cycle and dividing by the cycle time for that cycle. In the following sections, we use this data to answer two questions. Firstly, how large are the internal dissipations due to passive and motor friction relative to the external hydrodynamic dissipation? Secondly, what is the effect of the *Crisp2* gene deletion on flagellar energetics?

### D. Internal dissipation is larger than external hydrodynamic dissipation

The novel finding in Figure 4 (B) and (C) is that, for any WT or KO sample, the magnitudes of the internal frictional and motor dissipations are much larger than the dissipation in the external fluid. Before we examine this further, it must be reiterated that the results in Figure 4 for these dissipation rates depend on the values of the material parameters κ_0_ and *η*_0_. As previously mentioned, we have used here κ_0_ = 7 × 10^4^ Pa *μ*m^4^ based on experimental measurements elsewhere [9]. While the existence of internal friction in the fluid-filled region around the axoneme is expected [21: 28], direct measurements of the value of *η*_0_ are not available. For the same sperm motion quantified by the tangent-angle C-POD, we have calculated the energetics with different values of *η*_0_, ranging from zero to values well above the scaling estimate of 10^3^ Pa s *μ*m^4^. For any value of *η*_0_, we robustly find negative domains in the active power distribution, *p*^a^. However, as Figure 5 (A) shows, for nine out of the ten sperm samples studied, the minimum value of the net motor power delivered in a mean cycle, 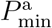, has a strongly negative value when *η*_0_ is much smaller than 10^3^ Pa s *μ*m^4^. In such a case, for a significant portion of the mean cycle around the time when 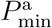 is negative, the axoneme as a whole does not drive the motion of the flagellum. The overall motion of the flagellum during that phase of the cycle is powered mostly by the release of the potential

**FIG. 5.**
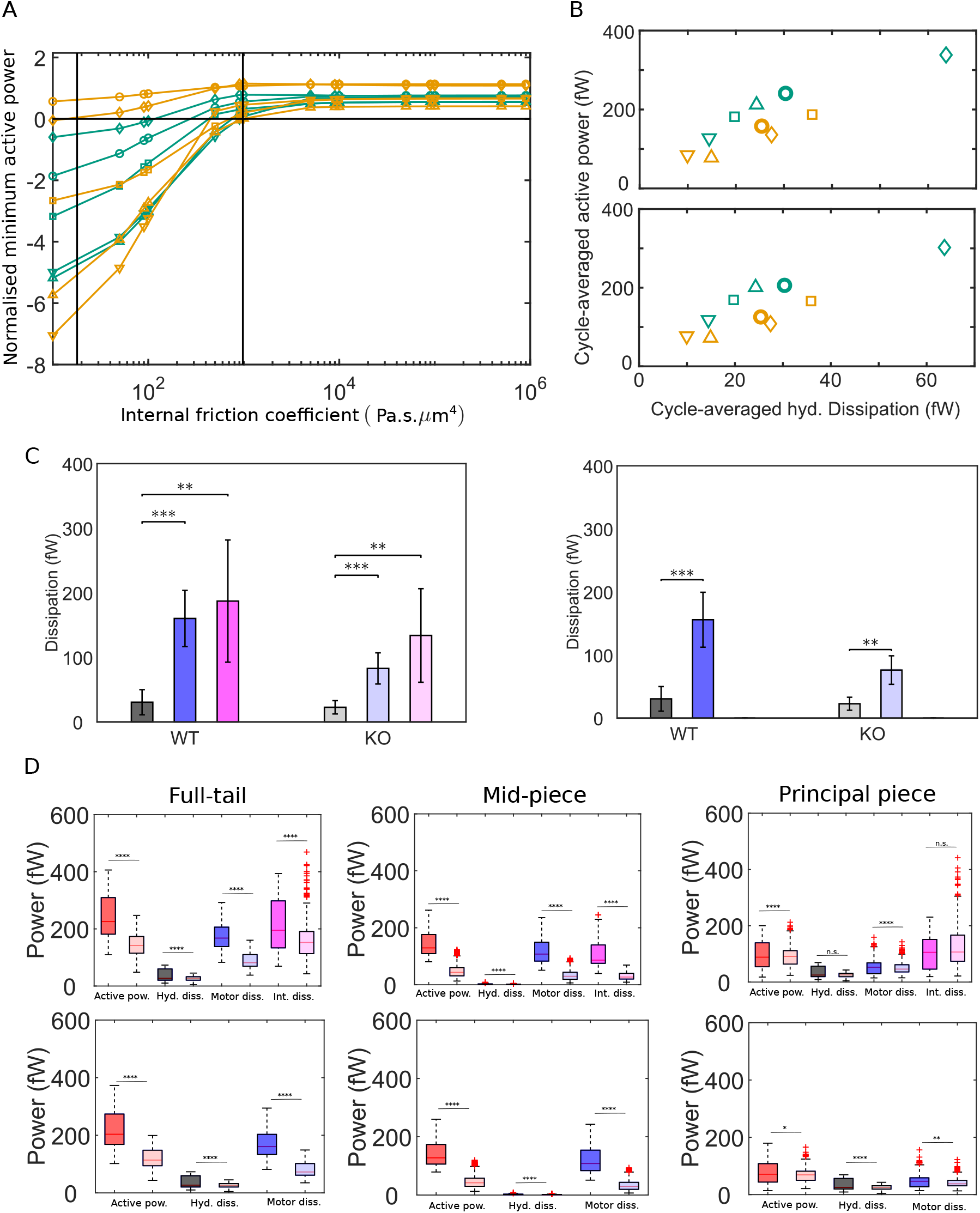
External versus internal dissipation in flagella. (A) Effect of the value of internal friction coefficient, *η*_0_, on the minimum value of the net active power required to overcome dissipation for WT (blue) and KO (red) samples. The minimum power value at each value of is normalised by the cycle-averaged active power at that across the mean cycle. The vertical line on the right is the scaling value of 10^3^ Pa s *μ*m^4^. It is also the minimum value of required for the net active power to always be positive at any phase of the mean cycle. The vertical line on the left is the value of energy stored elastically in the body of the flagellum. On the other hand, the value of 10^3^ Pa s *μ*m^4^is the value expected if the observed beat frequency of O(10) Hz were to be determined primarily by the interplay of filament elasticity and internal friction [21; 28], when external hydrodynamic friction is relatively negligible. Figure 5 (A) shows that, above this scaling estimate of *η*_0_, *P^a^_min_* is positive. The motion of the flagellum is strongly driven by the power input from the axoneme at all times during the beat cycle when *η*_0_ > 10^3^ Pa s *μ*m^4^.

**FIG. 6.**
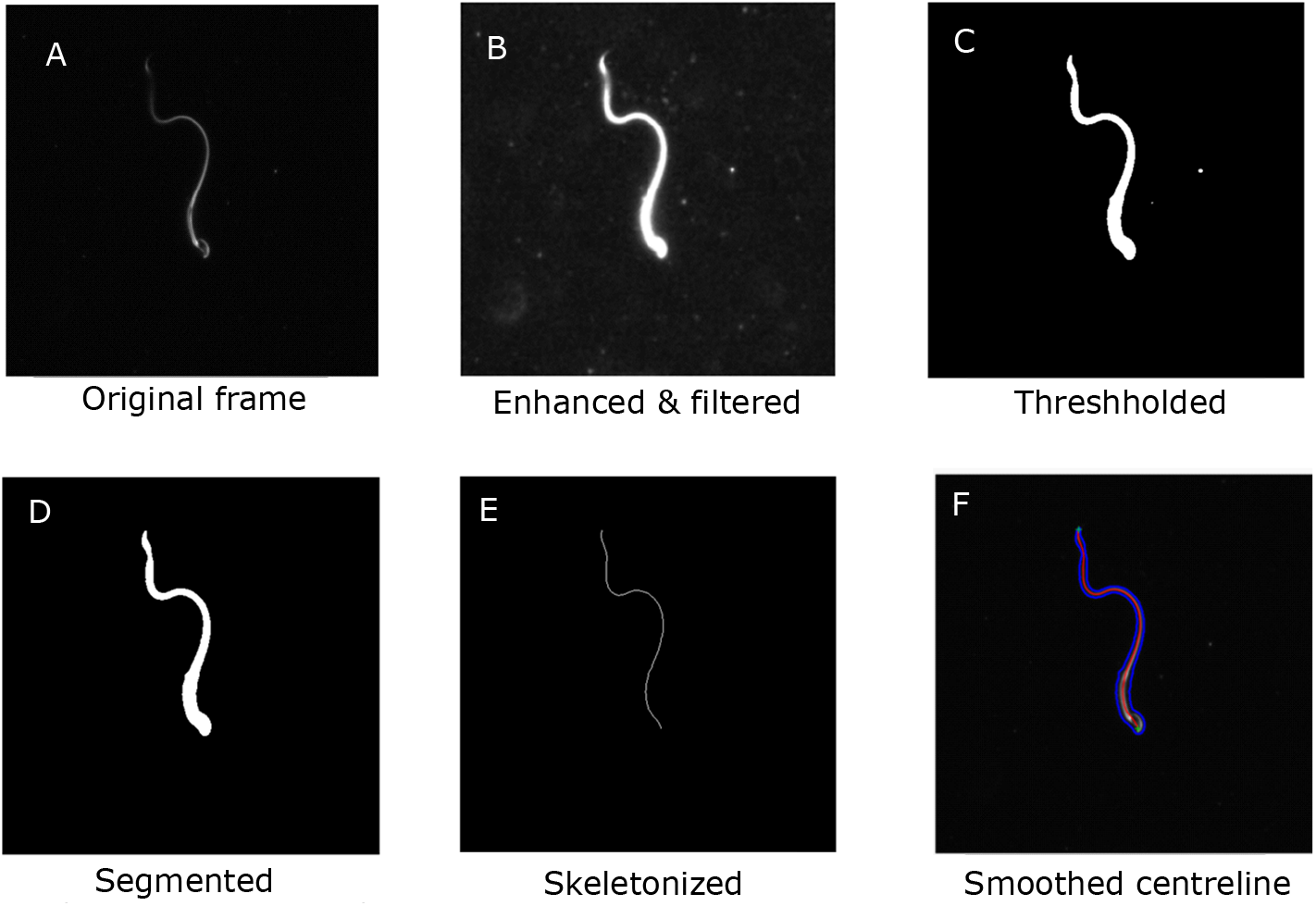
Main steps in the image-processing algorithm

In Figure 5 (B - D), we plot results for the energetic variables obtained with 10^3^ and *η*_0_ = 0 Pa s *μ*m^4^. With either value of *η*_0_, Figure 5 (B) shows that average net motor power input over a mean cycle for the five samples appears positively correlated with the cycle average of the hydrodynamic dissipation rate, within each genotype. The nature of the correlation depends on the genotype. Further, the excess of the cycle-averaged motor power input over the hydrodynamic dissipation is due to the internal dissipation. Considering the values for each sample in Figure 5 (B), we see that the internal dissipation must be significantly larger than the external hydrodynamic dissipation.

This is more clearly revealed in Figure 5 (C) where we have compared the population means of these different modes of dissipation. A one-way ANOVA reveals that, for all these quantities, the sample means in a genotype are distinctly different from the population mean for that genotype obtained by pooling all the cycles from the samples together (*P* ≪ 10^-4^; Table SI-1 in SI). Due to this large sample-to-sample variation in the sample means, the standard deviations in in Figure 5 (C) over the set of five sample means are large. Despite this, however, the differences between the external hydrodynamic dissipation on the one hand, and the internal motor and frictional dissipations on the other are sizeable enough to be statistically highly significant, as judged by unpaired, two-tailed Student *t*-tests (*p* ≪ 10^-4^; Table SI-2 in SI). When *η*_0_ = 10^3^ Pa s *μ*m^4^, in the WT samples, the motor (171 fW) and passive (210 fW) internal dissipations are 4.8 and 5.9 times larger than the hydrodynamic dissipation (35.4 fW), whereas in the KO samples, they are 3.4 (88.8 fW) and 6.2 (161 fW) times larger than the hydrodynamic dissipation (25.8 fW), respectively. Even when *η*_0_ is set to zero, the motor dissipations in the WT (166 fW) and the KO (80.4 fW) samples are 4.7 and 3.1 times larger than the hydrodynamic dissipation.

The differences in the energetics of the WT and KO populations are more subtle and the variation within the set of five sample means of each population is too large for drawing conclusions with confidence. We can, however, pool together all the individual cycles from each sample in a genotype to create a much larger dataset of individual time cycles for each genotype. The box-plots in Figure 5 (D) summarize the statistics of the entire pool of cycle-averaged powers for each genotype obtained with *η*_0_ = 10^3^ Pa s *μ*m^4^ (top-panel) and with zero internal friction (bottom-panel). Using Student t-tests with this large dataset (Table SI-3 in SI), we find that the net input from the dynein motors in sperm from *Crisp2* KO mice is significantly smaller than the power input in the corresponding WTs. This is observed over the entire tail. We further find that each kind of dissipation - hydrodynamic, motor or internal friction – is smaller in sperm from *Crisp2* KO mice. These observations in Figure 5 (D) are consistent with those in Figure 4 (A) that the *Crisp2* KO samples have smaller beating amplitudes over the entire flagellum. The rapidity of the beating, *i.e*. the mean beat frequency, is also an important factor in determining the rate of energy dissipation. In the samples studied here, due to the large variability in cycle-times, we do not find a significant difference (p > 0.01 in a Student t-test; Table SI-3) between the population means of the cycle-times (0.16 s and 0.18 s for WT and KO, respectively) or their reciprocals (7.19 Hz and 7.2 Hz, respectively) even after pooling the cycle-times from the samples from each genotype together. Investigations with a larger population size, however, reveal that the *Crisp2* KO does beat more slowly (7.4 Hz and 6.9 Hz in WT and KO, respectively; p = 0.05; [47]).

## III. DISCUSSION

The work presented here contributes to understanding flagellar beating in sperm on several fronts. We have shown that we can use conservation principles and take image-analysis of flagellar beating to extract, for the first time, the details of energy flows within sperm flagella. This ability to be able to study the energetics of live sperm is crucial to fundamentally understanding how the axoneme works and is regulated. We have further used the C-POD technique to unambiguously split the data into individual time cycles. This enables us to define mean cycles for all variables associated with flagellar beating. When used with tethered sperm, we can collect sufficient data to make statistically significant observations despite the large variability in beating patterns.

The data reveals several fascinating new insights into flagellar energetics. We firstly see that along the filament, there exist distinct phases during each cycle where dynein motors in the axoneme are driven back against the torques they exert by the motion of the rest of the flagellar body. It is known that dynein motors are regulated to create a travelling wave of forces, and hence turning moments, that propagates down the flagellum [59]. The periodic occurrence of positive and negative domains in the active power distribution in Figure 3 (D) shows that the local angular rate and the motor torque are in the same direction for some part of the beat cycle and in opposite directions in other parts of the beat cycle. In other words, the physical motion and the dynein forcing are out of phase with one another.

This implies that the work done on the motors in any region of the flagellum by the rest of the flagellar body during the negative-*p*^a^ phases of the beat cycle must be locally dissipated away quickly at that location. There is already evidence that dyneins can dissipate energy locally: it is known that dynein motors can cycle through conformational changes driven by ATP binding and hydrolysis even when not driving microtubule sliding [60]. Optical-tweezer experiments have further shown that dyneins can dissipate the work done on them when driven in the reverse by an external load by forces larger than the stall force for these motors [57]. Our results here reveal that this dissipation can be a dominant part of the energy budget within the flagellum and is likely to play an important role in determining the emergent dynamics of the filament. However, to our best knowledge, a mechanism of motor dissipation is not accounted for in models of axonemal driving [43].

In addition to motor dissipation, we find that the friction in the accessory structures surrounding the flagellum could also be significant. While most current models of flagella or cilia assume that the flagellum is a purely elastic filament, it is recognized that internal friction could play a role in flagella and cilia [28; 61]. No measurements of the internal friction coefficient in flagella are available. The results Figure 5 (C) show that, if the friction coefficient is large enough to determine the beating frequency of the flagellum [21; 28], the magnitude of the internal dissipation caused by this friction is considerably greater than the hydrodynamic dissipation in the aqueous medium outside. Development of systematic techniques to independently characterize the visco-elastic characteristics of the non-axonemal part of the flagellum is necessary if we are to understand the physics of flagellar beating through comparing model predictions with experiment.

Our experiments were conducted with an aqueous buffer. If the kinematics of the beating pattern remain unchanged, when fluid inertia is negligible, a higher medium viscosity would result in a proportionately higher hydrodynamic dissipation. While the internal frictional dissipation would remain unchanged, the magnitudes of the motor power distribution, and hence the motor dissipation, would also increase. However, it is known that the kinematics of beating dramatically changes with an increase in medium viscosity [62]. It is therefore difficult to directly extrapolate the results here to predict how the relative magnitudes of the internal and external dissipations would change in a more viscous medium.

We have shown that the passive friction and motor dissipations are features that are shared across sperm from the WT and the *Crisp2* KO genotypes. The CRISPs are the subclade of the CAP superfamily proteins that are expressed in the male reproductive tract. Although CRISPs are not essential for fertility [33; 48; 63], we see here that a lack of CRISP2 significantly reduces the mechanical power input from the axoneme in sperm, which in turn appears to be responsible for slower beating with smaller amplitude. CRISP2 is further known to be incorporated internally into the sperm flagellum [64] and is expected to act by regulating ion channels on the cell or organelle membranes [33; 47]. The approach presented here can similarly be used to systematically explore the role played by other proteins and signalling agents on the regulation of flagellar beating.

## IV. MATERIALS AND METHODS

### A. Sperm sample preparation

Generation of knockout mouse models and all animal procedures were approved by the Monash University Animal Experimentation Ethics Committee. The mouse knockout line were maintained on a C57/BL6N background. Sperm were collected from *cauda epididymides* and *vas deferens* using the back-flushing method [33] in modified TYH medium (135 mM NaCl, 4.8 mM KCl, 2 mM CaCl_2_, 1.2 mM KH_2_PO_4_, 1 mM MgSO_4_, 5.6 mM glucose, 0.5 mM Na-pyruvate, 10 mM L-lactate, 10 mM HEPES, pH7.4). The samples were stored in dark at 37 deg C until imaging. Sperm samples WT-1 and -2 were from the same indvidual, WT-3 and -4 were from another individual, and WT-5 was from a third individual mouse. All the five KO samples were from separate individuals.

### B. Tethering and imaging

Sperm motility was investigated in a custom-made observation chamber. Briefly, two strips of double-sided tape (90 *μ*m nominal thickness) were affixed to a glass slide 16 mm apart. A drop of 40 *μ*l of sperm suspension was placed between the two strips and sealed against evaporation with 17 mm square coverslips (Thermo Fisher Scientific, No. 1.5). The TYH medium was supplemented with 0.3 mg/ml of BSA to cause cells to adhere to the glass slide. Only sperm tethered at their heads with freely beating flagella were chosen for video imaging and subsequent analysis.

An Olympus AX-70 upright microscope equipped with a U-DFA 18 mm internal diameter dark-field annulus, an 20x 0.7 NA objective (UPlanAPO, Olympus, Japan) and incandescent illumination served as the platform for the imaging system. All extraneous optical elements were removed from the detection light paths to maximise system light efficiency. An ORCA-Flash4.0 v2+ (C11440-22CU) sCMOS camera (Hamamatsu, Japan) was used for capturing images. This system leverages a high frame-rate for motion capture, an exceptional 82% QE for the low level of light and the small 6.5 *μ*m pixel size to increase system spatial resolution [65–67].

The optical lateral resolution was 0.479 *μ*m at a reference wavelength of 550 nm. With the 6.5 *μ*m pixel size of the ORCA sCMOS and the system magnification factor of 20, the best-case lateral resolution of 0.650 *μ*m (325 *μ*m/pixel) at the Nyquist-Shannon sampling was sufficient to spatially resolve the tip of the sperm tail. A 512 x 512 pixel region of interest therefore corresponded to an experimental sample field of view (FOV) of 166.4 x 166.4 *μ*m, which was sufficient for most of the experiments reported here. Occasionally, sperm with stiffer flagella, required an FOV increase with a reduction of approximately 0.8 fps for each pixel increase.

Image data was free-streamed to a Xeon E5-2667 computer (with a 12-core CPU running at 2.9 GHz supplemented by 64 GB of DDR3 RAM and 1 TB SSD hard drive in a RAID0 configuration) via a dedicated Firebird PCIe3 bus 1xCLD Camera Link frame-grabber card (Active Silicon, United Kingdom) at the 8.389 MB/s memory buffer speed of the camera. This resulted in a capture frame rate of approximately 400 frames per second (fps). The best-case blur-free motion capture of the system at this frame rate corresponds to element point velocities of 130 *μ*m/s. The Fiji image-processing package was used for image capture control along with the Micro-Manger Studio plugin (version 1.4.23) for multi-dimensional acquisition [66] set to 4000 time points, zero time point interval, a 2.0 ms exposure time. The data was written as an image stack.

Camera resolution can be increased to exceed optical resolution by replacing the 180 mm tube lens with a 250 mm tube lens. The region of interest would then increase to 714 x 714 pixels, with the capture frame rate being reduced to approximately 286 FPS. Frame exposure can be likewise increased to 3.25 ms to allow for a superior signal to noise ratio (SNR).

### C. Image-analysis and skeletonization

The mean of the grayscale intensity at each pixel location across all the frames was used to construct a background image. This was then subtracted from each frame to remove the background. The contrast was then adjusted to enhance the foreground grayscale intensity. Median filters of different sizes were applied to remove noise. The grayscale image was then smoothened with a Gaussian filter before binarization at a threshold computed by Otsu’s method [68]. Connected components in the binarized image were then located and classified according to size and eccentricity. The sperm body is expected to have the largest size among the objects in the frame. An oval (i.e. an ellipse) is fitted around each body. The eccentricity is a measure of the deviation of the oval from a perfect circle. An oval fitted around the whole sperm body will be highly elongated and will have a high eccentricity. These two criteria were used to automatically identify the sperm body in each frame and remove other extraneous objects. Morphological thinning was then applied to the segmented image to extract a skeleton of the sperm tail. Spurious branches on the skeleton were automatically identified and removed to give an unbranched skeleton. The skeleton at this stage is rough, with noisy burrs that are then smoothed out using low-pass filtering. The resulting smoothed curve representing the sperm body is henceforth referred to as the centerline. Since the algorithm processes each frame independently of all others, the processing can be run in parallel. 1000 imaged frames divided into sets of 200 frames each were processed in parallel on a high-performance computing cluster.

The arc-length between each adjacent pair of points was calculated and the overall contour length of the centerline in each frame was obtained. Motion of the sperm body out of the plane of focus leads to blurring and loss of contrast and intensity of the image, which in turn, increases errors in the automated processing of the images. This is particularly problematic at the tail end of the flagellum. As a result, the skeleton obtained is truncated at the tail end, resulting in a loss of total contour length of the captured skeleton. Videos with significant loss of length were discarded, and only videos showing largely in-plane beating, with deviations smaller than 10% from the mean segmented length were considered for further analysis.

For each video, the time-averaged end-to-end straight line was first determined. Sperm centerlines in every frame were rotated by an angle to align this line with the horizontal *x*-axis. Mouse sperm have hook-shaped heads. The centerlines in a video were reflected about the horizontal axis if necessary, to orient the head-hook concave downwards in all videos. At this stage, the pixel points on the centerline were not uniformly distributed along the length of the sperm body. That is, the arc length between each adjacent pair of points is not the same along the centerline. The *x* and *y* coordinates for each centerline point were linearly interpolated to obtain a large number of points (~ 200) distributed uniformly with the same difference in the arc-length *s* between adjacent points. Frames were also not always equally spaced in time since poor-quality frames were discarded. Linear interpolation in time was applied across the two frames on either side of a missing frame to compute the centerline in the missing frame. Tangent angles to the horizontal were computed at each s in every frame. A Butterworth low-pass filter was used to spatially smoothen the tangent-angle-versus-s data in each frame.

### D. Data processing

There are four stages in calculating flagella energetics, starting from the raw tangent-angle data obtained from the centerlines after image-processing. This original tangent-angle function is denoted as 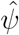.

1. Firstly, at each time instant, a 20th-order Chebyshev polynomial is fitted through the pixel data in the tail region such that it is adequately smooth across the neck. The boundary condition at the junction is obtained from the data in the head region after ensuring that the rigid-head condition is met. The combined intermediate tangent-angle profile for the rigid head and the Chebyshev-polynomial tail is denoted as 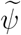.
2. The C-POD is performed to obtain an optimally compact representation of the tail region in the form shown in Equation 17.
3. The tangent-angle profile thus obtained, *ψ*(*s, t*), is used to calculate other geometric, kinematic and dynamic quantities.
4. Mean cycles of all physical quantities and the standard errors in the means are then calculated.

In the description below, we denote time-averages, 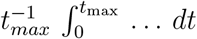 as 〈…〉. The imaged tail region is defined as *s* ∈ [*s*_N_, *s*_T_], where *s*_T_ is the maximum value of *s* for which pixel data is available for every time sample (typically, *s*_T_ = 0.85 *L*). In this region, we work with a rescaled variable that maps the domain [*s*_N_, *s*_T_] onto [-1,1]:

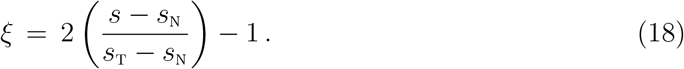

The inner-product of a pair of functions *f* and *g* with respect to a weighting function *w*(*ξ*) is defined as 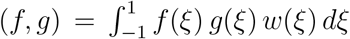 with the norm of *f* given by 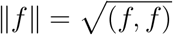.

#### 1. Intermediate tangent-angle profile

A Chebyshev polynomial,

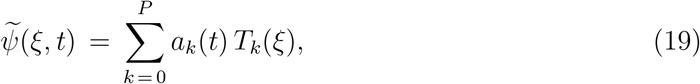

of order *P* = 20 is fitted to the data in the imaged tail region at each time, *t*. Here, *T_k_* is the *k*-th Chebyshev polynomial of the first kind [69]. To ensure *C*^2^-continuity across the neck, the Chebyshev polynomial must satisfy the following boundary conditions at *ξ* = −1, the neck:

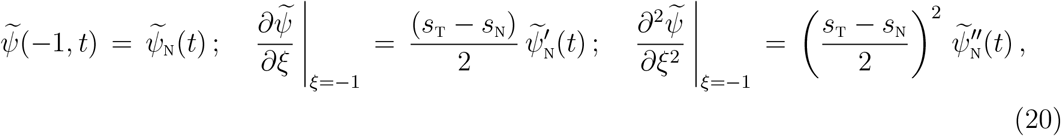

where 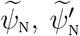 and 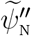 are the values of the tangent angle and its first two *s*-derivatives at the neck, respectively. We discuss the calculation of these values for the rigid head shortly.

Using the properties of Chebyshev polynomials and Lagrange’s method of undetermined coefficients, we determine the set of coefficients *a_k_* that minimize 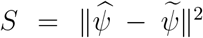, the mean square-error between the raw data, 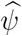, and the Chebyshev polynomial, 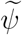, weighted by 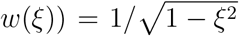, while also satisfying the constraints specified by Equation 20 Using standard methods and Gaussian quadrature to approximate the integrals, we obtain

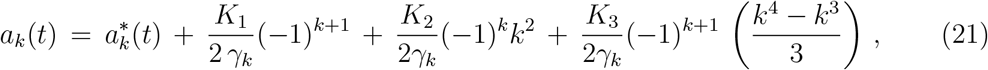

where γ_*k*_ = (1 + *δ*_0, *k*_)/(2 (*P* + 1)), *δ_i,j_* is the Kronecker *δ*-function, *K*_1_, *K*_2_ and *K*_3_ are the Lagrange multipliers. The unconstrained Chebyshev coefficient,

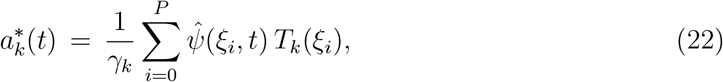

where *ξ_i_* is the *i*-th root of the *P* + 1-th Chebyshev polynomial [54]. The values of *T_k_* (*ξ_i_*) can be calculated using standard recursion relations [54]. Substituting from Equation 21 in the boundary conditions into Equation 20 results in a system of linear equations that can be solved for the Lagrange multipliers. Inserting these values back into Equation 21 gives the Chebyshev coefficients in the imaged tail region. The resulting 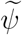 is consistent with boundary conditions at the neck.

The raw centerline data 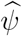 in the head region, *s* ∈ [0, *s*_N_), does not satisfy the rigid-head condition, primarily because the image of the head is large and diffuse, leading to errors in identifying its centerline consistently. To impose the rigid-head conditions, *∂C/∂t = ∂ω/∂s* = 0, the time-averaged tangent-angle profile 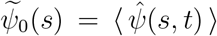 is first calculated in this domain. Then, the tangent-angle profile in this domain is set to the following to ensure the rigid-head conditions:

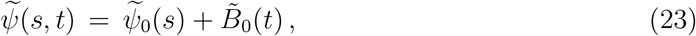

where,

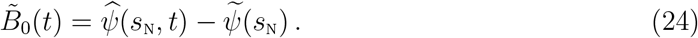

The time average of *B*_0_ is thus zero. With this profile, the values 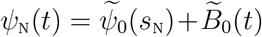. The time-independent *s*-derivatives 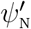 and 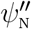 are determined from the time-averaged data points from values of 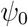 adjacent to the neck using second-order backward-difference formulae. The rotation rate of the whole head-region, 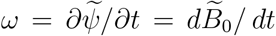, is non-zero.

#### 2. C-POD of the tail region

The C-POD provides advantages over the ‘empirical’ POD used previously for sperm [34; 35]. The empirical POD is applied directly on the discrete data to produce shape modes that are numeric vectors. The discrete nature of the modes makes high-order spatial derivatives computed from them susceptible to noise. The C-POD approach here allows derivatives to be computed without noisy artefacts. Further, specific restrictions on the shape at the boundaries can be conveniently imposed.

We first recall key aspects of the general POD technique to obtain the optimal mutually orthogonal basis functions. The Chebyshev polynomials *T_k_* themselves constitute a set of mutually orthogonal basis functions. At any *t*, 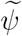 is a polynomial of order P that is expanded in terms *P* + 1 Chebyshev polynomials. Given a small number *M < P* + 1, say *M* = 2, any linear combination of *M* of the Chebyshev polynomials can be expected to be a poor approximation of the full *P*-th order polynomial, 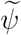. The technique of POD allows us to find a set, {*ω_m_*}, of *M* unique orthogonal functions different from *T_k_* such that a linear combination of these provides the *best* approximation of 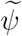 possible, given the choice of *M*. The gain is that we need to track only the set of *M* coefficients {*B_m_*} as functions of time rather than the larger set of all the *P* + 1 time-dependent Chebyshev coefficients, {*a_k_*}.

The time-averaged profile in the imaged-tail region is 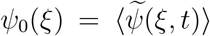. The deviation of the original function 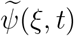 from this time-average is

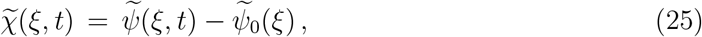

and the spatial two-point cross-correlation of 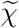 is

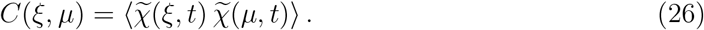

It can be shown that the set of optimal basis functions for the POD are the eigenfunctions of this two-point cross-correlation [36; 37]. That is, an optimal shape mode, *ω_m_*, is such that

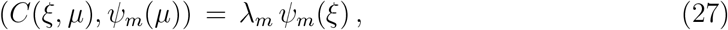

where λ_*m*_ > 0 is the corresponding eigenvalue. These eigenfunctions are mutually orthogonal *i.e*. (*ω_m_, ω_n_*) = *δ_m,n_*. The coefficient of the *m*-th shape mode is then obtained by projection as

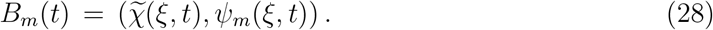

These coefficients are themselves orthogonal in time, i.e., 〈*B_m_*(*t*) *B_n_*(*t*)) = *δ_m,n_ λ_m_*.

The matrix algorithm for obtaining the time-independent Chebyshev coefficients of the shape modes is as follows. The Chebyshev polynomials are first normalized as follows:

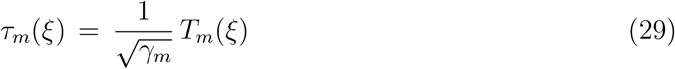

so that the inner-product (with the Chebyshev weighting function) (*τ_m_,τ_n_*) = *δ_m,n_*. The Chebyshev coefficients *a_k_* of 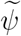 are correspondingly rescaled as 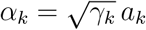, so that 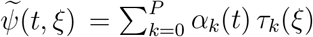. The Chebyshev coefficients of the time-averaged tangent-angle profile, *ω*_0_(*ξ*), and the deviation from the mean, 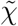 are then 〈*α_k_*(*t*)) and Δ*α_k_*(*t*) = *α_k_*(*t*) − (*a_k_*(*t*)), respectively. From Equation 27, the cross-correlation, 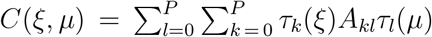, where

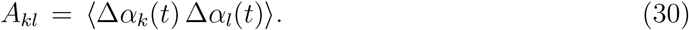

The symmetric matrix A composed of A_*kl*_ is equivalent to the cross-correlation matrix. Diagonalizing the matrix A = V · Λ · V^T^ yields the *P* +1 eigenvalues, {λ_*m*_}, of the correlation operator as the diagonal elements of the matrix, Λ. The *m*-th column of V is the m-th eigenvector of A. Its elements are the Chebyshev coefficients of the *m*-th shape mode:

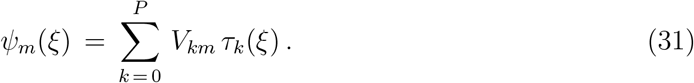

The corresponding shape coefficient can be obtained from the equation above and from Equation 28 as

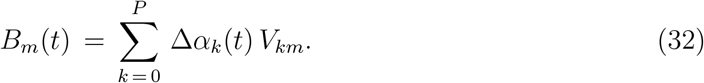

With *ψ_m_*, and *B_m_* thus determined from the original cross-correlation of 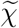, we can obtain the C-POD approximation, *ψ*, given by Equation 17 for any choice of *M* ≤ *P* + 1. The deviation of the C-POD approximation from the mean,

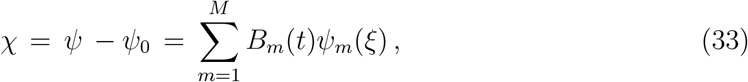

is an approximation of the original 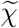. The approximation improves with increasing *M* and when 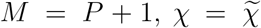 exactly, since the full set of *P* + 1 eigenfunctions {*ψ_m_*} spans the same function space that is spanned by the set of P + 1 Chebyshev polynomials, {*T_k_*}. Further, using the orthogonality of the shape modes, it can be shown that

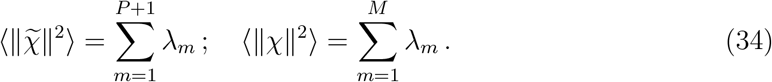

Therefore, the mean-squared error in the approximation when *M* < *P* + 1,

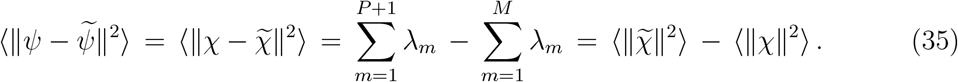

We can, therefore, use the ratio of the cumulative sum of the eigenvalues for any *M*, normalized by the sum of all the *P* + 1 eigenvalues,

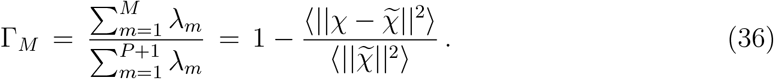

as a measure of the accuracy of the *M*-th order C-POD representation: the closer Γ_*M*_ is to 1, the better *ω* captures 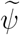. As discussed earlier, 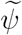 is constructed to be consistent with the neck boundary conditions (in Equation 20) at all times. The C-POD basis functions, 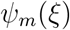, that span this function space, therefore, also satisfy the same neck boundary conditions.

#### 3. Calculation of flagellar kinematics and dynamics

The equations in are used to calculate the active power distribution in the following manner:

1. The centerline coordinates are obtained from the tangent angle *ψ*(*s, t*) by:

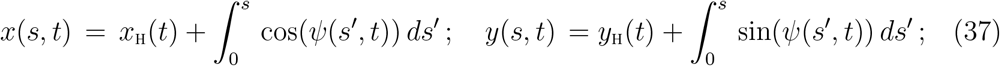

where *x*_H_ and *y*_H_ are the experimentally-determined coordinates of the tip of the head at any time *t*. Further, (*t_x_, t_y_*) = (cos *ω*, sin *ω*) and (*n_x_, n_y_*) = (−sin *ψ*, cos *ψ*).
2. Since the shape modes are given by Equation 31, their spatial derivatives are calculated by applying standard recursion relations for the Chebyshev polynomials [54]. The time rates of the shape coefficients, 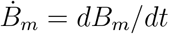, are calculated numerically using central-difference formulae.
3. We then calculate the spatial derivatives of the C-POD approximant, *ψ* and obtain the curvature, *C = ∂ψ/∂s*, and its derivatives. Its time derivative, 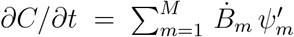 and the centerline angular velocity, 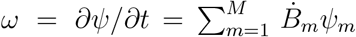 are calculated.
4. Noting that,

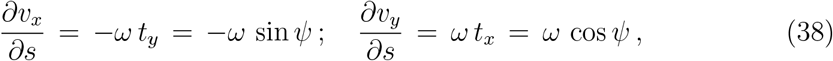

flagellar velocities are calculated from the rotation rate as follows:

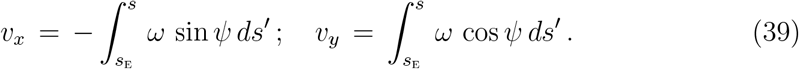

where *s*_E_ is the experimentally-determined location of the tether point. The tangential and normal components of the centerline velocity, *υ_t_* = v · t and *υ_n_* = v · n, are then calculated.
5. The hydrodynamic force distribution, f^h^, is calculated using Equation 10 and the expressions for the tangential and normal friction coefficients.
6. Equation 3 together with Equation 10 for the hydrodynamic force f^h^ gives

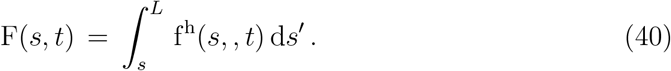
7. M^*E*^, M^*D*^, and *ϵ* are calculated using the constitutive equations Equation 12 and Equation 13; M = M^*E*^ + M^*D*^.
8. The elastic stiffness and internal dissipation profiles, *κ*(*s*) and *η*(*s*), are obtained using Equation 14 with the chosen values of *κ*_0_ and *η*_0_.
9. The distributions 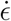 and *p*^i^ are calculated using Equation 15 *p*^h^ is calculated using v and f^h^ in Equation 6, *p*^s^ is calculated with using Equation 7.
10. [Equation 16 is used to obtain the active power distribution, *p*^a^.
11. Since the head is modeled as a rigid, passive, body, 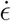, *p*^i^ and *p*^a^ are zero in that region. The total instantaneous power dissipated by the head against the external hydrodynamic and tethering forces,

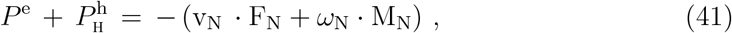

the power delivered on to head by the force F_N_ and moment *ω*_N_ acting on the neck junction.

#### 4. Mean beat cycles

The time-dependent coefficients of the dominant shape modes, *B*_1_ and *B*_2_, are plotted against one another. Individual beat cycles are identified from the times at which the polar angle of a point in this *B*_1_–*B*_2_ space is zero. In other words, a beat cycle starts when the flagellar shape is a scaled version of the first shape mode, *ψ*_1_. The time-phase within the *i*-th beat cycle is then calculated as

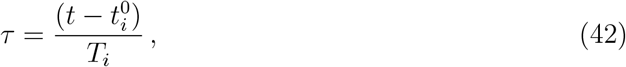

where 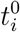 is the starting time of the *i*-th cycle and 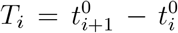 is the time-period of that cycle. Any function of the tangent angle and its derivatives with respect to either *s* or *t* can be split into individual beat cycles and can be expressed as functions of the time-phase, *τ*. The mean cycle of that function over the set of its cycles can be computed at each *τ* and so can the standard-error in the mean (s.e.m.) Between 40 and 60 beat cycles were captured for each sperm sample.

Ref. [70] was used to obtain the red-blue colormap for the power kymographs. Ref. [71] was used to plot the shaded error bars on the kymographs.

## V. ACKNOWLEDGMENTS

This research was funded by the Australian Research Council (DP190100343, DP200100659), an Interdisciplinary Research Grant from the Provost’s office at Monash University, IITB-Monash University Academy funding, and the Department of Biotechnology within the Government of India (# BT/PR13442/MED/32/440/2015).

## VI. AUTHOR CONTRIBUTIONS

R. P., S. J. and A. N. designed the research, developed the model equations and the analytical tools, and drafted the paper. The experiments were designed and performed by M.K.O.B, A.G. and D. P. All authors were involved in data analysis and interpretation and editing the manuscript.

## I. CONSERVATION LAWS

The position vector of any material point on a cross-section is = r + R, where r is the point on the cross-section through which the filament axis passes and R is the vector displacement of the material point from the axial point. Then, the velocity of the material point is 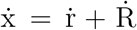. Cross-sectional planes can rotate relative to each other. Then, *∂* d_*i*_/∂*t* = *ω* × d_*i*_, where *ω*(*s, t*) is the instantaneous angular velocity of the crosssectional plane through the axial point at *s*. The velocity of any material point, *∂*x/*∂t* = v + *∂*R/*∂t*, where v = *∂*r/*∂*t. With this, we have *∂*R/*∂t* = *ω* × R. It can further be shown that,

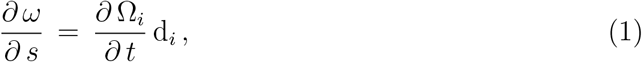

where Einstein’s summation convention is used. This implies that, for planar motion, where *ω* = *ω* e_*z*_,

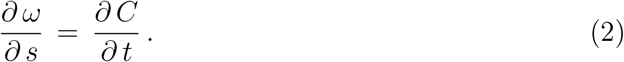

Using kinematic relationships and the Frenet-Serret equations, it can be shown that

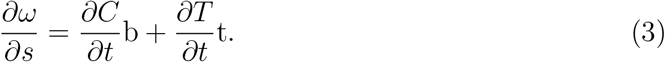

Mass conservation is trivially satisfied for rods whose density is constant and crosssectional area is independent of time. The net hydrodynamic force on a cross section,

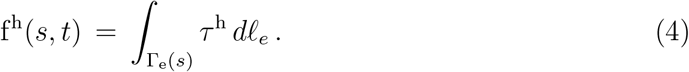

Here, *τ*^h^ is the hydrodynamic traction acting on the external surface and Γ_e_(*s*) is the external perimeter of the cross-section at any *s*, parameterised by an arc-length variable along the perimeter, *ℓ*_e_. The external force distribution f^e^(*s, t*) similarly accounts for non-hydrodynamic surface traction such as that due to wall contact. The axonemal motors exert forces on the *internal* surfaces of the passive flagellar material. The surface traction, *τ*^a^, exerted by these motors results in an active force distribution,

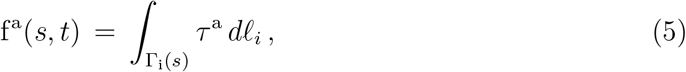

where Γ_i_(*s*) is the perimeter at any section *s* of the internal surfaces and *ℓ_i_* is an arc-length variable along that perimeter. The passive material stress tensor *σ* acting throughout the cross-sectional domain ∑(*s*) results in a net force, F, on a cross-section by the material on its aft side:

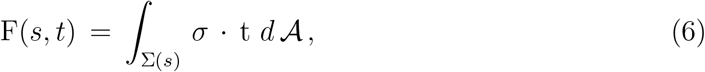

where 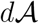 is a differential area element on a cross-section at *s*. The torques due to the hydrodynamic and motor tractions are

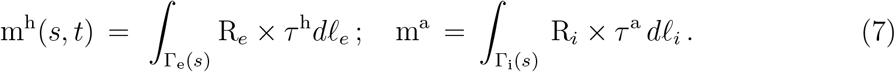

The torque distribution, m^e^, due to the external non-hydrodynamic surface traction is similarly defined. The torque due exerted by the material on the aft side,

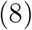

The conservation of linear momentum for a section of the rod from *s*_1_ to *s*_2_ is given by:

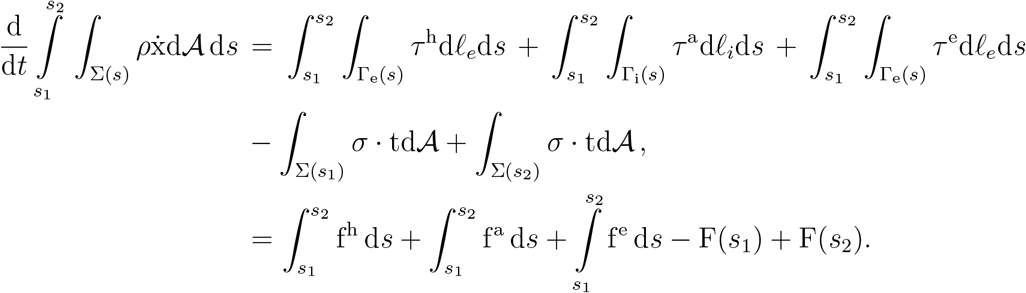

For an inertialess rod, the differential form of the equation above is

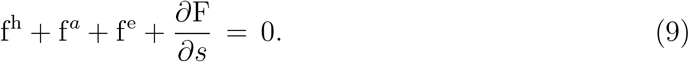

The gradient with respect to s of F in the momentum balance thus describes the net force per unit length at a cross-section due to the passive internal stress.

Similarly, the conservation of angular momentum for a section of the rod can be written as follows,

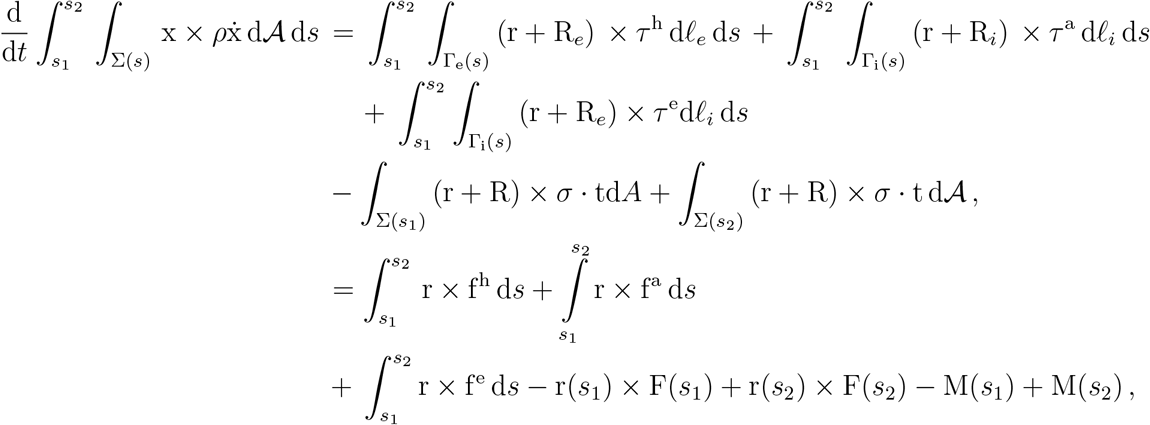

which leads to the following differential equation for an inertialess rod after eliminating terms using the differential form of the linear momentum equation earlier and t = *∂*r/*∂*s:

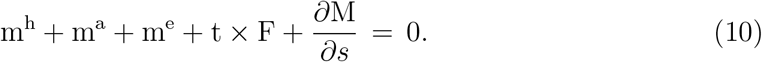

The First Law of Thermodynamics provides an equation that balances the rate of energy change with the work done and heat input to a control volume. In the case of an internally-driven rod, the total energy is the sum of the passive elastic energy, the thermal internal energy and the kinetic energy. Work is done on the control volume by the surface tractions exerted by the surrounding fluid, the internal motors and the external tethering constraint. Work is also done by the passive material stress on the cross sections. Heat can transferred out of the control volume to the surroundings. The conservation of energy implies:

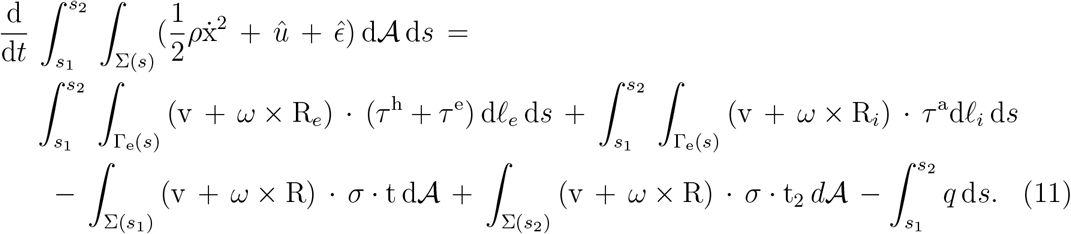

Here, 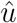 and 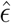 are the thermal internal energy and elastic strain energy densities, and *q* is the heat transferred per unit length out of the cross-section at *s*. Defining the energy distributions 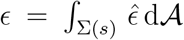 and 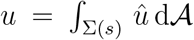, and neglecting kinetic energy changes in an inertialess rod, the differential form of the energy equation is obtained as

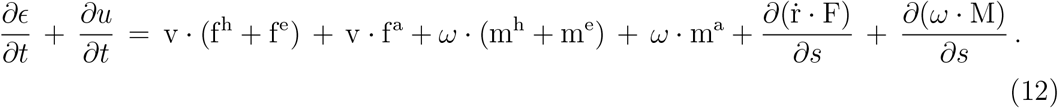

## II. STANDARD RECURSION RELATIONS FOR CHEBYSHEV POLYNOMIALS

The Chebyshev polynomials of the first kind are defined by a recursive relation of the following form:

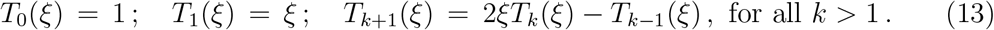

The first and second derivatives of the polynomials are then derived as:

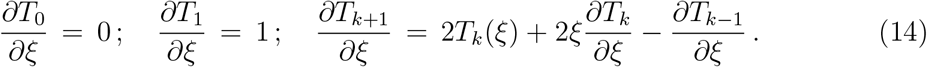

Further, at *ξ* = −1, *T_k_* = (−1)^*k*^ and *∂T_k_/∂ξ* = *k*^2^(−1)^*k*−1^.

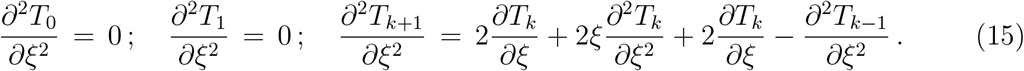

The roots of the *P* + 1-th Chebyshev polynomial,

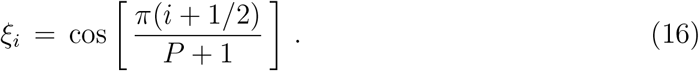

## III. SUPPLEMENTARY RESULTS

**FIG. 1:**
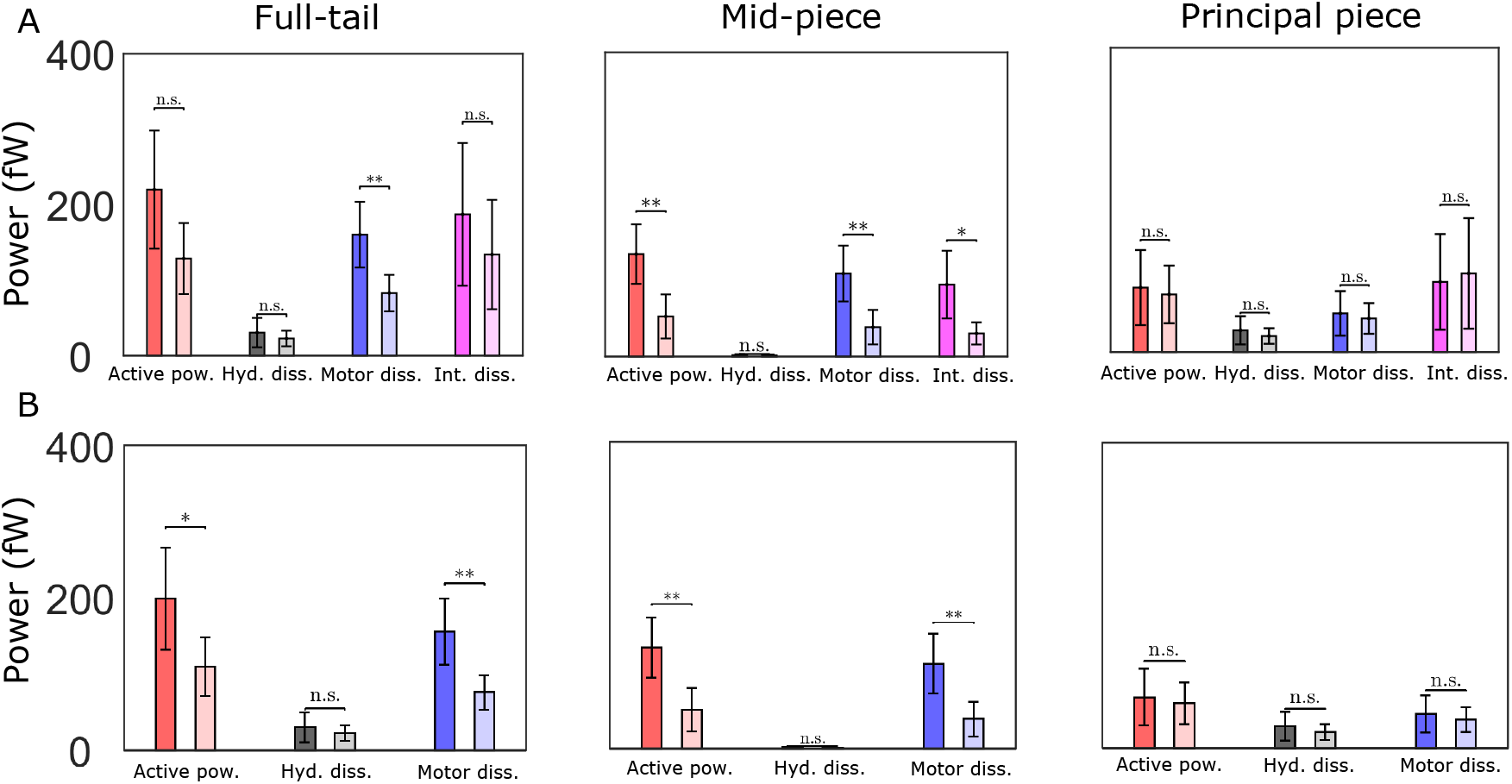
Cycle-averaged powers. The data shown are population means and variances obtained with the sample-means from five WT and CRISP2 KO sperm. Unpaired two-tailed t-tests are used to compare population means; ** to refers *p* ≤ 10^-3^, * to *p* ≤ 0.05. Differences are not significant (n.s.) when *p* > 0.05.

**TABLE I:**
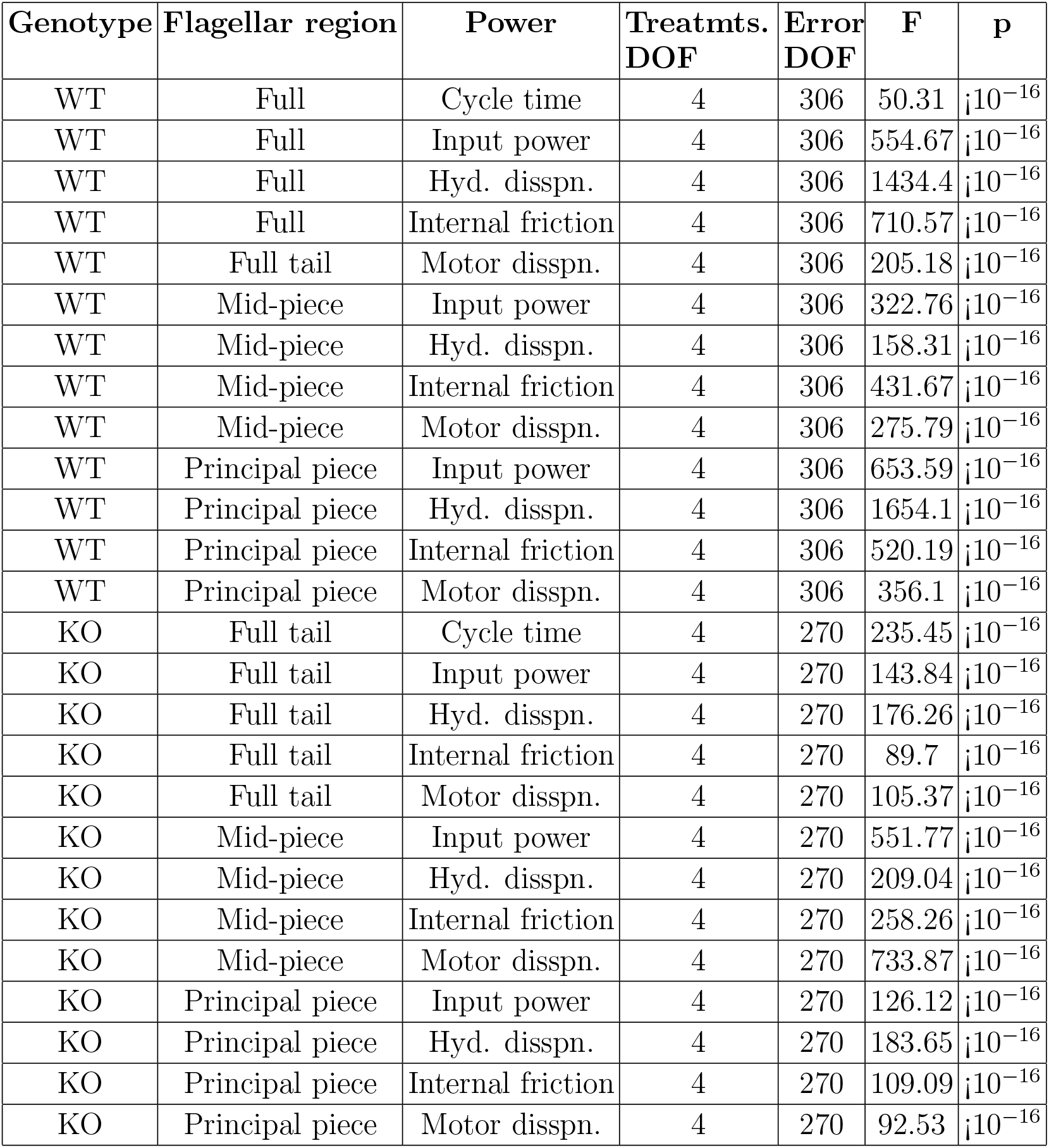
One-way ANOVA data for establishing that sample means of cycle variables are significantly different from population means in the WT and KO genotypes.

**TABLE II:** Comparison of the external hydrodynamic dissipation with the internal frictional and motor dissipations

| Genotype | Flagellar region | $\eta_0$ Pa s $\mu\text{m}^4$ | (Hyd. disspn(fW)) | Int. friction (fW) | t | p |
| --- | --- | --- | --- | --- | --- | --- |
| WT | Full tail | $10^3$ | 35.40 | 209.80 | 32.88 | $< 10^{-16}$ |
| WT | Mid-piece | $10^3$ | 2.14 | 105.50 | 38.37 | $< 10^{-16}$ |
| WT | Principal piece | $10^3$ | 33.21 | 103.82 | 19.95 | $< 10^{-16}$ |
| KO | Full tail | $10^3$ | 25.79 | 160.53 | 28.06 | $< 10^{-16}$ |
| KO | Mid-piece | $10^3$ | 1.49 | 31.51 | 32.65 | $< 10^{-16}$ |
| KO | Principal piece | $10^3$ | 24.26 | 128.82 | 21.28 | $< 10^{-16}$ |
| Genotype | Flagellar region | $\eta_0$ Pa s $\mu\text{m}^4$ | Hyd. disspn(fW) | Motor. disspn. (fW) | t | p |
| WT | Full tail | $10^3$ | 35.40 | 170.86 | 47.8 | $< 10^{-16}$ |
| WT | Mid-piece | $10^3$ | 2.14 | 115.63 | 50.78 | $< 10^{-16}$ |
| WT | Principal piece | $10^3$ | 33.21 | 54.90 | 11.33 | $< 10^{-16}$ |
| KO | Full tail | $10^3$ | 25.79 | 88.76 | 38.94 | $< 10^{-16}$ |
| KO | Mid-piece | $10^3$ | 1.49 | 37.26 | 25.34 | $< 10^{-16}$ |
| KO | Principal piece | $10^3$ | 24.26 | 51.36 | 18.24 | $< 10^{-16}$ |
| WT | Full tail | 0 | 35.40 | 166.44 | 45.78 | $< 10^{-16}$ |
| WT | Mid-piece | 0 | 2.14 | 118.12 | 49.26 | $< 10^{-16}$ |
| WT | Principal piece | 0 | 33.21 | 47.96 | 8.56 | $< 10^{-16}$ |
| KO | Full tail | 0 | 25.79 | 80.41 | 35.57 | $< 10^{-16}$ |
| KO | Full tail | 0 | 1.49 | 37.42 | 25.44 | $< 10^{-16}$ |
| KO | Principal piece | 0 | 24.26 | 42.87 | 14.72 | $< 10^{-16}$ |

**TABLE III:** Comparison of WT and KO samples. The values in in columns 2 and 3 are the means of the distributions created by pooling together values from the individual cycles of all the sperm samples in each genotype.

| Quantity | WT | KO | t | p |
| --- | --- | --- | --- | --- |
| Cycle time (s) | 0.16 | 0.18 | -2.72 | 0.01 |
| Reciprocal cycle time (Hz) | 7.19 | 7.2 | -0.03501 | 0.97 |
| Full tail hyd. dissipation (fW) | 35.40 | 25.79 | -7.39 | $5.17 \times 10^{-13}$ |
| Mid-piece hyd. dissipation (fW) | 2.14 | 1.49 | -7.59 | $1.26 \times 10^{-13}$ |
| Principal piece hyd. dissipation (fW) | 33.21 | 24.26 | -7.19 | $2 \times 10^{-12}$ |
| Full tail motor dissipation (fw), $\eta_0 = 10^3 \text{ Pa s } \mu\text{m}^4$ | 170.86 | 88.76 | -26.36 | $< 10^{-16}$ |
| Full tail int. friction (fw), $\eta_0 = 10^3 \text{ Pa s } \mu\text{m}^4$ | 209.80 | 160.53 | -6.93 | $1.11 \times 10^{-11}$ |
| Mid-piece motor dissipation (fw), $\eta_0 = 10^3 \text{ Pa s } \mu\text{m}^4$ | 115.63 | 37.26 | -28.80 | $< 10^{-16}$ |
| Mid-piece int. friction (fw), $\eta_0 = 10^3 \text{ Pa s } \mu\text{m}^4$ | 105.50 | 31.51 | -24.74 | $< 10^{-16}$ |
| Principal piece motor dissipation (fw), $\eta_0 = 10^3 \text{ Pa s Pa s } \mu\text{m}^4$ | 54.90 | 51.36 | -1.66 | 0.098 |
| Principal piece int. friction (fw), $\eta_0 = 10^3 \text{ Pa s Pa s } \mu\text{m}^4$ | 103.82 | 128.82 | 4.29 | $2.1 \times 10^{-5}$ |
| Full tail motor friction (fw), $\eta_0 = 10^3 \text{ Pa s Pa s } \mu\text{m}^4$ | 166.44 | 80.41 | -27.69 | $< 10^{-16}$ |
| Mid-piece motor dissipation (fw), $\eta_0 = 10^3 \text{ Pa s Pa s } \mu\text{m}^4$ | 118.12 | 37.42 | -28.49 | $< 10^{-16}$ |
| Principal piece motor dissipation (fw), $\eta_0 = 10^3 \text{ Pa s } \mu\text{m}^4$ | 47.96 | 42.87 | -2.83 | 0.0048 |

